# A peptide of a type I toxin-antitoxin system induces *Helicobacter pylori* morphological transformation from spiral-shape to coccoids

**DOI:** 10.1101/585380

**Authors:** Lamya El Mortaji, Alejandro Tejada-Arranz, Aline Rifflet, Ivo G Boneca, Gérard Pehau-Arnaudet, J. Pablo Radicella, Stéphanie Marsin, Hilde De Reuse

## Abstract

Toxin-antitoxin systems are found in many bacterial chromosomes and plasmids with roles ranging from plasmid stabilization to biofilm formation and persistence. In these systems, the expression/activity of the toxin is counteracted by an antitoxin, which in type I systems is an antisense-RNA. While the regulatory mechanisms of these systems are mostly well-defined, the toxins’ biological activity and expression conditions are less understood. Here, these questions were investigated for a type I toxin-antitoxin system (AapA1-IsoA1) expressed from the chromosome of the human pathogen *Helicobacter pylori*. We show that expression of the AapA1 toxin in *H. pylori* causes growth arrest associated with rapid morphological transformation from spiral-shaped bacteria to round coccoid cells. Coccoids are observed in patients and during *in vitro* growth as a response to different stress conditions. The AapA1 toxin, first molecular effector of coccoids to be identified, targets *H. pylori* inner membrane without disrupting it, as visualized by Cryo-EM. The peptidoglycan composition of coccoids is modified with respect to spiral bacteria. No major changes in membrane potential or ATP concentration result from AapA1 expression, suggesting coccoid viability. Single-cell live microscopy tracking the shape conversion suggests a possible association of this process with cell elongation/division interference. Oxidative stress induces coccoid formation and is associated with repression of the antitoxin promoter and enhanced processing of its transcript, leading to an imbalance in favor of AapA1 toxin expression.

Our data support the hypothesis of viable coccoids with characteristics of dormant bacteria that might be important in *H. pylori* infections refractory to treatment.

**Significance Statement:** *Helicobacter pylori*, a gastric pathogen causing 800,000 deaths in the world annually, is encountered both *in vitro* and in patients as spiral-shaped bacteria and as round cells named coccoids. We discovered that the toxin from a chromosomal type I toxin-antitoxin system is targeting *H. pylori* membrane and acting as an effector of *H. pylori* morphological conversion to coccoids. We showed that these round cells maintain their membrane integrity and metabolism, strongly suggesting that they are viable dormant bacteria. Oxidative stress was identified as a signal inducing toxin expression and coccoid formation. Our findings reveal new insights into a form of dormancy of this bacterium that might be associated with *H. pylori* infections refractory to treatment.

## Introduction

Toxin-antitoxin (TA) systems are small genetic elements that are widely distributed on bacterial mobile genetic elements and among archaeal and bacterial genomes, often in multiple copies (1–4). They code for a small stable toxic protein, designated toxin, whose action or expression is counteracted by an unstable antitoxin molecule that can either be an RNA or a protein (3, 5). Under conditions in which the toxins can act, they target essential cellular processes or components (transcription, DNA replication, translation, cell wall or membrane) resulting in growth arrest or cell death (3, 5). The molecular mechanism of the toxins’ activities and their regulation are often known in detail while the biological function remains elusive. TA systems take part in plasmid maintenance and protection from phage infection. Less is known on the biological functions and targets of the chromosomally encoded TA systems. Some are important for bacterial survival in their mammalian host (6) or biofilm formation (7). There is accumulating evidence that upon stress conditions, some TA systems play a role in the switch of actively growing bacteria to persisters or dormant cells (8–11). Persister bacteria constitute a subpopulation that is metabolically active but slow-growing and highly tolerant to antibiotics and stress conditions (11). Persistence is considered as an important cause of recalcitrance of chronic bacterial infections to therapy. However, stress-induced activation of TA-encoded toxins does not necessarily cause persister formation (12) and it was more generally found that decreased intracellular ATP concentration and/or low proton motive force (PMF) are landmarks of persister formation (13, 14).

In the present work, we explored the role of TA systems in *Helicobacter pylori*, a bacterium that colonizes the stomach of half of the human population worldwide and causes the development of gastritis. In some cases, gastritis evolves into peptic ulcer disease or gastric carcinoma that causes about 800,000 deaths in the world every year (15, 16). This microaerophilic bacterium is unique in its capacity to persistently colonize the stomach despite its extreme acidity and intense immune response (16). The molecular mechanisms at the origin of this exceptional adaptation capacity of *H. pylori* remain only partially understood. Their elucidation is crucial to understand *H. pylori* virulence and to improve its eradication in patients with recurrent peptic ulcer disease. *H. pylori* is a Gram-negative bacterium with a helical shape. Upon stress conditions (antibiotics, aerobic growth) or prolonged culture, the shape of *H. pylori* progressively evolves into a U-shape followed by a spherical form designated coccoid (17, 18). *H. pylori* coccoids are non-culturable bacteria proposed to be dormant forms (18). Recently, dormant non-culturable cells were associated with a deeper state of dormancy as compared to persister cells (19).

*H. pylori* coccoids were observed in human gastric biopsies and, like spirals, adhere to gastric epithelial cells (20). Despite numerous reports on coccoid forms of *H. pylori*, no cellular effector of this conversion has been reported so far.

In type I TAs, the antitoxin is a small regulatory RNA inhibiting the synthesis of the toxin by base pairing the toxin-encoding mRNA (3–5, 21, 22). Four families (A-B-C-D) of conserved type I TA systems are expressed from the chromosome of *H. pylori*, only the A family was studied as it is highly expressed and conserved among *H. pylori* strains (23–25). For the A1 TA system, the detailed mechanism by which transcription of the IsoA1 antitoxin RNA impairs AapA1 toxin synthesis by base pairing with its primary transcript, ensuring both translation inhibition of the AapA1 active message and leading to rapid degradation of the duplex by RNase III, has been recently published (24, 25). The *H. pylori* type I toxins are typically small hydrophobic peptides of 30-40 amino acids predicted to form alpha-helices (26). No clues on the mode of action or the physiological role of these systems have been reported.

Here we show that the AapA1 toxin induces a rapid morphological transformation of *H. pylori* from spirals to coccoids by targeting the inner membrane and probably interfering with cell elongation and division. Furthermore, oxidative stress induces coccoid formation and triggers imbalanced expression of the TA components in favor of toxin production.

## Results

### The AapA1 toxic peptide, expressed by the AapA1/IsoA1 TA system, triggers rapid transformation of *H. pylori* into coccoids

Recently, the study of the AapA1/IsoA1 TA system of *H. pylori* (**Fig. 1A**) showed that expression of the AapA1 toxin (a 30 amino acids-long hydrophobic peptide) leads to bacterial growth arrest (24). Given the genetic organization of the type I AapA1/IsoA1 locus in B128 strain, we decided to investigate the mechanism underlying the activity of AapA1 by using a strain in which AapA1 is under the control of an inducible promoter. We thus transformed *H. pylori* strain B128 deleted of the AapA1/IsoA1 chromosomal locus with each of the three plasmids constructed by Arnion *et al*. (24) (**Fig. 1A**). The first, pA1-IsoA1, derived of the pILL2157 vector (27) contains a functional TA locus in which the AapA1 gene is under the control of an IPTG inducible promoter (24). In plasmid pA1, derived from this first construct, the antitoxin IsoA1 promoter has been inactivated by point mutations that do not interfere with transcription or translation of the AapA1 ORF (24). Derived from this second construct, plasmid pA1* contains an additional mutation that inactivates the start codon of the AapA1 peptide (24). Under our experimental conditions, we observed, in agreement with data of (24), that addition of IPTG did not significantly influence the growth rate of strains harboring pA1-IsoA1 or pA1*. In contrast, addition of the inducer causes a rapid growth arrest of *H. pylori* with pA1 indicating a toxic effect of AapA1 expression (**Fig. 1B**). The growth arrest was accompanied by loss of culturability of more than 10^3^-fold 8h after induction.

**Figure 1:**
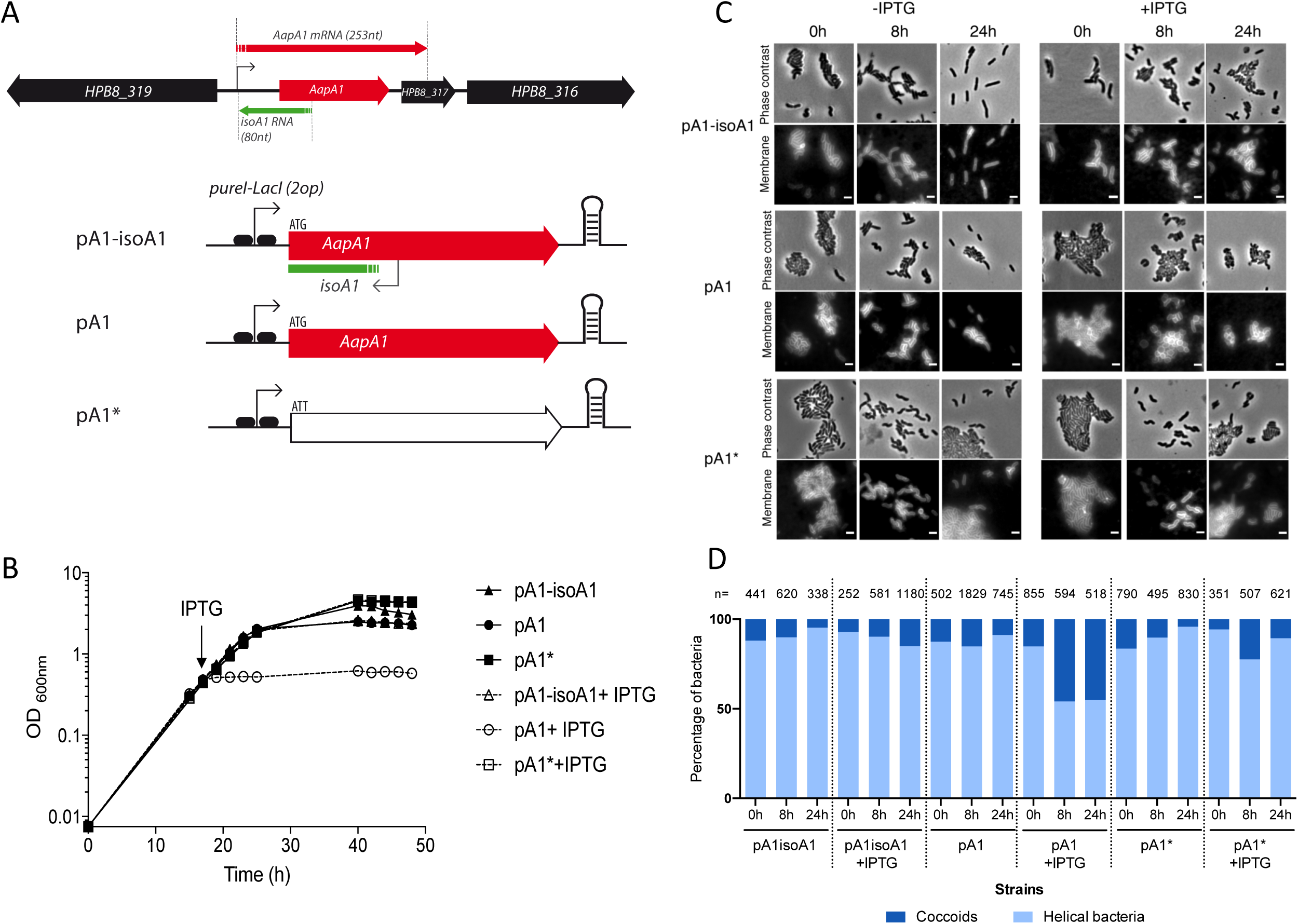
Expression of the AapA1 toxin from the AapA1/IsoA1 TA system induces growth arrest and transformation of *H. pylori* strain B128 into coccoids. **A) Genetic organization of AapA1/IsoA1 locus in *H. pylori* B128 strain with the AapA1 transcript encoding the toxin and the *IsoA1* antitoxin RNA**. Below is the representation of the inserts of the three plasmids used in this study (24), each derived from the pILL2157 *E. coli/H. pylori* vector (27). These inserts are expressed under the control of an IPTG inducible promoter. Plasmid pA1-IsoA1 expresses both the toxin and the first 30 nucleotides of the *IsoA1* RNA antitoxin. In pA1, the promoter region of IsoA1 was mutated without affecting the amino acid sequence of AapA1. Plasmid pA1* was derived from pA1 and contained an additional mutation inactivating the start codon of AapA1. **B) Expression of the AapA1 toxin causes rapid growth arrest of *H. pylori***. Growth kinetics of *H. pylori* B128 strains carrying each of the three plasmids illustrated in panel A, grown in the absence (black symbols) or in the presence of 1mM IPTG (empty symbols) that was added at the time indicated by an arrow (16h). **C) Expression of the AapA1 toxin induces transformation of *H. pylori* into coccoids**. Microscopy analysis of the *H. pylori* B128 strains carrying each of the three plasmids at 8h and 24h following IPTG addition or grown without IPTG. Phase-contrast and fluorescence images are respectively represented on the upper and bottom panels. *H. pylori* membranes were stained with the lipophilic dye FM4-64. The scale bar corresponds to 2 μm. **D) Quantification of the transformation of *H. pylori* into coccoids**. The proportion of helical and coccoid bacteria under each condition was quantified with the phase contrast images using the MicrobeJ software. The number of analyzed cells is indicated for each condition.

In parallel to growth, we investigated the consequences of AapA1 expression on *H. pylori* morphology. Samples from bacterial cultures with these plasmids were grown in the presence or absence of IPTG, stained with a membrane-specific dye, FM4-64, and analyzed by fluorescence microscopy at 0h, 8h and 24h (**Fig. 1C**). *H. pylori* cells expressing the wild-type AapA1/IsoA1 locus (pA1-IsoA1) or the pA1* mutated locus present a classical helical rod-shaped phenotype upon IPTG exposure (**Fig. 1C**). Under the same conditions, we observed that cells carrying the pA1 plasmid, and thus expressing the AapA1 toxin exhibit a rapid morphological conversion to spherical cells also known as coccoid forms (**Fig. 1C**). Using MicrobeJ (28) analysis of the phase contrast images, we quantified the relative proportions of coccoids vs spiral bacteria under each condition (**Fig. 1D**). When the toxin is expressed (pA1+IPTG), about half of the cells have converted into coccoid as soon as 8h post-induction without further changes at 24h post-induction. No bacterial lysis was detected even at 24h post-induction. With strains carrying the pA1* plasmid, significant amounts of coccoids were only observed after 64h of culture, similarly to the control wild type strain containing an empty plasmid. These data show for the first time that the AapA1 toxin induces a fast conversion of *H. pylori* cells from spiral-shaped to coccoid forms.

### Inner membrane targeting by the AapA1 toxin

To investigate the mode of action of the AapA1 toxin, we first analyzed its subcellular localization in *H. pylori* using a GFP reporter. A C-terminal fusion of the toxin with GFP (A1-GFP) was introduced into B128 strain deleted of the chromosomal AapA1/IsoA1 module and expressed either from the chromosome under control of its native promoter or from the IPTG-inducible promoter of vector pILL2157 (27) (**Fig. S1**). After IPTG induction, the plasmidic fusion did not affect *H. pylori* growth indicating that a C-terminal GFP fusion prevents the toxicity of the toxin. Attempts to construct N-terminal tagged-AapA1 were unsuccessful.

Cellular fractionation was then performed on the strains, and the subcellular localization of the fused toxin was analyzed by immunoblotting. **Figure 2A** shows that the AapA1-GFP fusions expressed from the chromosome or from the plasmid are almost exclusively present at the inner membrane of *H. pylori*. No A1-GFP peptide was detected in the culture medium. Fluorescence microscopy of live bacteria after induction of the AapA1-GFP fusion revealed a weak patchy pattern in the periphery of the cell (**Fig. 2B**), different from a GFP-alone control cell (**Fig. S2**), and compatible with membrane localization. This observation was confirmed by measuring the overlap of the fluorescence intensity profiles of the toxin and TMA-DPH membrane dye measured perpendicular to the length axis of *H. pylori* (**Fig. 2B**).

**Figure 2:**
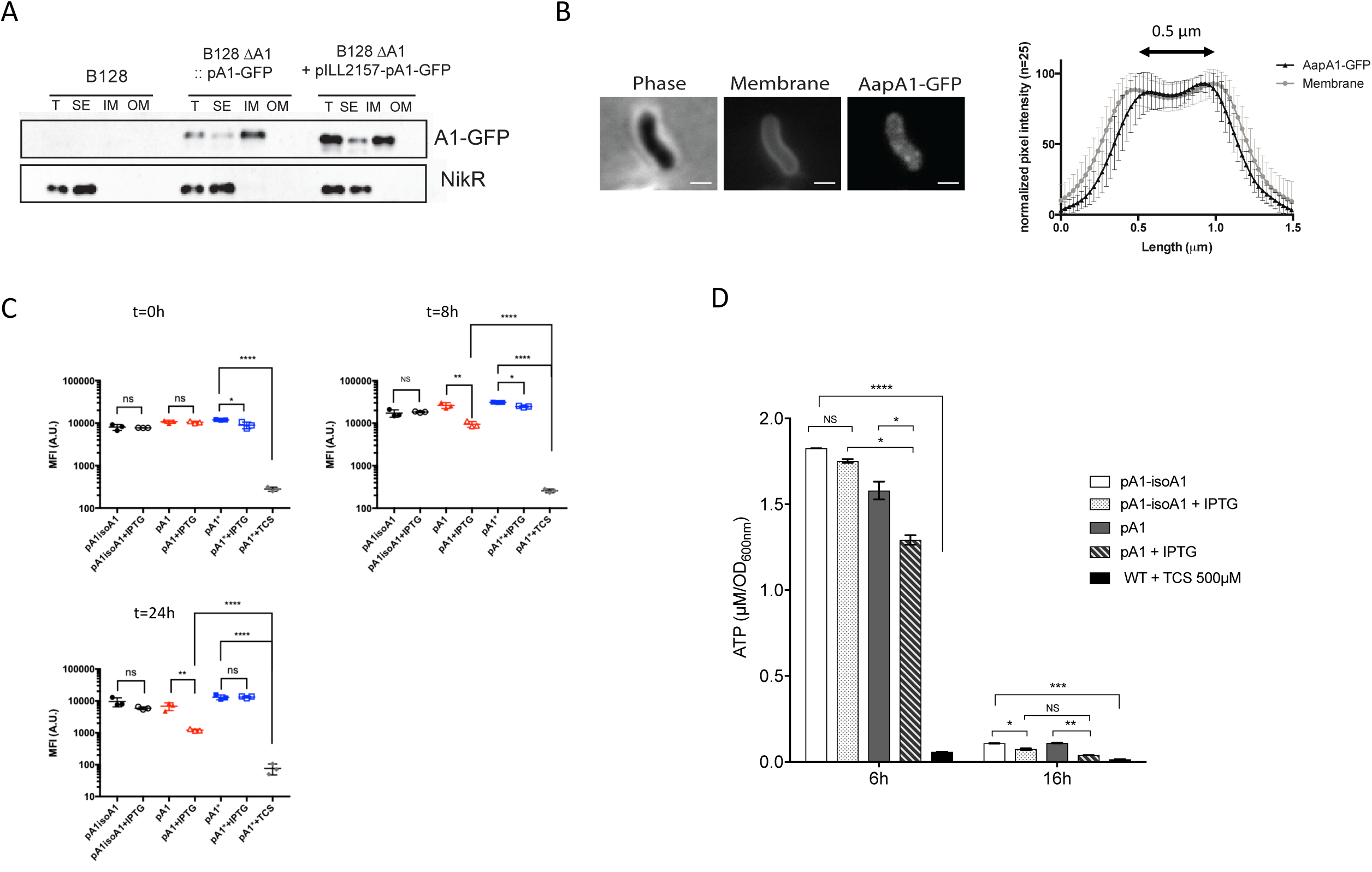
AapA1 toxin targets *H. pylori* inner membrane, weakly disturbs the bacterial membrane potential and poorly affects intracellular ATP content. **A) AapA1 toxin localizes to the *H. pylori* inner membrane**. Western blot analysis of total extracts (T), soluble extracts (SE), inner membrane (IM) and outer membrane (OM) fractions prepared from *H. pylori* B128 strain expressing a GFP tagged-AapA1 toxin (pA1-GFP) expressed either from the chromosome or from a plasmid (pILL2157) under the control of an IPTG-inducible promoter. Anti-NikR antibodies were used as a control of the cytoplasmic fraction. The fractionation procedure was validated as shown in **Fig. S2**. **B) In live cells, AapA1-GFP localizes as discrete foci at the *H. pylori* membrane**. Strain B128 expressing the AapA1-GPF fusion protein was analyzed on agarose pads by fluorescence microscopy 6h after IPTG-induction. Membranes were stained with TMA-DPH lipophilic dye. The membrane association of AapA1-GPF was quantified by measuring the fluorescence intensity profile perpendicular to the length axis of *H. pylori*. The graph shows the average fluorescence intensity profiles with standard deviation (n=25). The fluorescence maxima separated by 0.5 μm correlate with the *H. pylori* cell width. **C) Analysis of the effects of the AapA1 toxin on *H. pylori* membrane potential**. MitoTracker Red CMXRos, a membrane potential reactive dye was used to analyze live *H. pylori* B128 strains expressing each of the three plasmids illustrated in **Fig. 1**. Samples were taken at 0, 8 and 24h post-induction, stained with MitoTracker Red CMXRos and the proton-motive force (PMF) was measured as the Mean Fluorescence Intensity (MFI) by flow cytometry. The cell population distribution histograms are presented in **Fig. S3**. Cells expressing the toxin (pA1) present a mild fluorescence reduction after 8 or 24h of culture suggesting a weak disturbance of their membrane potential. No changes were observed in cells containing either pA1-isoA1 or pA1* plasmid. As a control, pA1*-bearing cells were treated with TCS (3,3’,4’,5-Tetrachlorosalicylanilide), a protonophore active on *H. pylori*, resulting in a massive loss of membrane potential. The experiment was performed three times. Error bars represent the standard deviation, with * corresponding to *P* < 0.05, ** to *P* < 0.01, *** to *P* < 0.001; **** to *P* < 0.0001, indicating that the mean values are significantly different, and NS corresponding to non-significant, (*P* >0.05). **D) Measurement of intracellular ATP content**. Intracellular ATP was extracted from B128 strains harboring pA1-IsoA1 or pA1 at 6 or 16h post-induction growth in the presence or absence of IPTG. B128 WT strain treated with 500μM of TCS protonophore was used as a control of PMF dissipation. ATP concentrations were determined using a luciferase-based assay (BacTiter-Glo™, Promega). Results from 3 independent experiments performed in triplicates are shown. Error bars represent the standard deviation, with * corresponding to *P* < 0.05, ** to *P* < 0.01, *** to *P* < 0.001, **** to *P* < 0.0001, indicating that the mean values are significantly different, and NS corresponding to non-significant, (*P* >0.05).

From these results, we conclude that the AapA1 toxin is specifically targeted to the inner membrane of *H. pylori*.

### AapA1 toxin has a minor impact on *H. pylori* membrane potential and intracellular ATP content

Considering the membrane localization of the toxin, we hypothesized that AapA1 expression could affect the membrane potential and by consequence the ATP content of the cells. To explore this possibility, *H. pylori* cells harboring pA1-IsoA1, pA1* or pA1 plasmids were taken at different culture time points (0, 8 and 24h), exposed to MitoTracker Red CMXRos, a red-fluorescent dye whose accumulation inside live bacteria depends upon membrane potential, and fluorescence of individual cells was measured by flow cytometry. As a control, fluorescence was measured for the pA1* strain treated with 3,5,3’,4’-tetrachlorosalicylanilide (TCS), an effective *H. pylori* protonophore (29). The TCS treatment causes a massive drop in fluorescence due to dissipation of membrane potential (**Fig. 2C, Fig. S3**). We observed that, when the AapA1 toxin is expressed in *H. pylori*, the membrane potential is maintained. Only a weak drop was observed at 6h after induction that was slightly more pronounced at 24h (**Fig. 2C**).

Next, the ATP content of *H. pylori* strains carrying either pA1-IsoA1 or pA1 was measured from total metabolites extracted at different culture time points and quantified by a luciferase based-assay (**Fig. 2D**). After 6h of culture, IPTG induction had a minor effect on cellular ATP content of both strains as compared to the drastic consequences observed with the TCS control. At 6h post-induction, many pA1-containing cells have already transformed into coccoids (see live imaging data below). After 16h post-induction, in stationary phase, the overall ATP content strongly drops in both strains and IPTG induction causes an additional weak decrease that is poorly changed upon toxin expression by pA1 (**Fig. 2D)**.

Taken together, these results show that the AapA1 toxin slightly perturbs the *H. pylori* membrane potential with no major consequences on the cellular ATP content. We conclude that dissipation of membrane potential is not the major cause of toxin-induced bacterial growth arrest.

### Ultrastructural analysis of toxin-induced coccoids

For the first time, cryo-electron microscopy (cryo-EM) was used to visualize *H. pylori* coccoids in near-native states. Using this method, we compared exponentially growing bacteria, toxin-induced coccoids and “aging coccoids” (70h growth) (**Fig. 3**). In exponential phase, the characteristic helicoidal shape of *H. pylori* (strains B128 WT, 4h and pA1-IsoA1, 4-8h) was perfectly visible with a uniformly contrasted cytoplasm surrounded by two dense layers of membranes. The periplasm of variable thickness surrounding the cell is distinguishable as the low-density space between the inner and outer membrane. Strains expressing the toxin (pA1 at 4-8h) have intact flagella and are visible in two major morphotypes. In the first “U” shaped morphology, a round intact outer membrane surrounds a bent intact inner membrane. The second morphotype corresponds to round coccoids of a diameter of approximately 1µm, with visible integrity of both membranes and a central uniformly stained dense cytoplasm. This second morphotype is similar to that of aging coccoids from the WT strain (70h).

**Figure 3:**
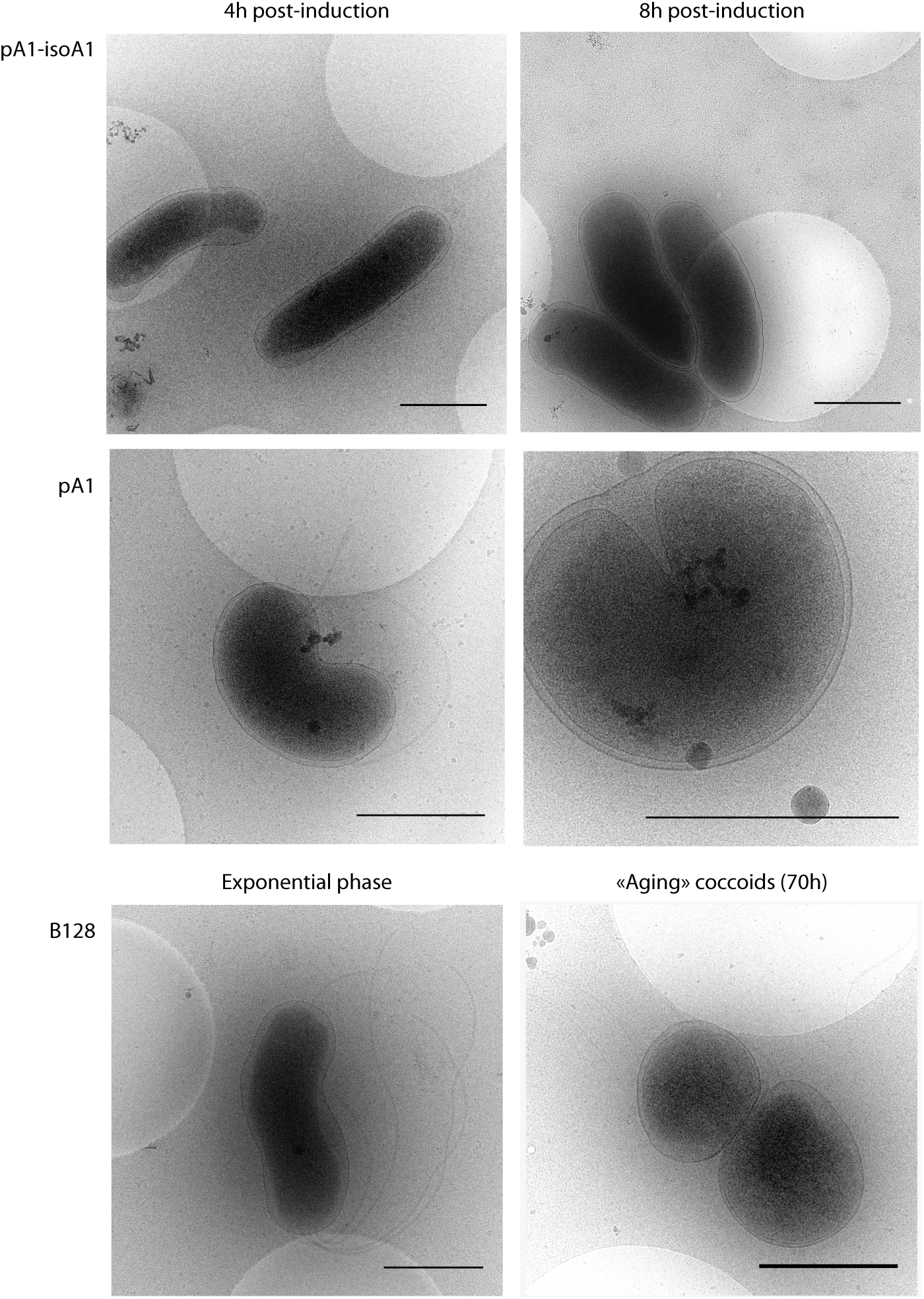
Ultrastructural analysis of toxin-induced coccoids and “aging” coccoids by cryo-electron microscopy. Cryo-electron microscopy was used to compare “aging” coccoids (70 h-old cultures) and exponentially growing B128 strains either WT or carrying plasmid pA1-IsoA1 or pA1 grown for 4 or 8h in the presence of IPTG. Expression of the toxin results in the formation of U-shaped bacteria at 4h and round coccoids cells at 8h with visible intact bacterial cell envelope similar to the aging coccoids. Scale bar represents 1μm.

In conclusion, cryo-EM images revealed that the toxin-induced *H. pylori* coccoids have no visible defect in membrane integrity and are ultrastructurally similar to “aging coccoid” forms.

### Modification of peptidoglycan composition upon transformation of *H. pylori* into coccoids

Our data showed that the AapA1 toxin targets *H. pylori* inner membrane without causing significant membrane potential collapse and without visible loss of integrity. Therefore, we tested whether the toxin-induced cell shape transformation could be associated with modifications in the cell wall and thus in peptidoglycan (PG) composition. PG extracted from B128 WT strain in exponential (24h of culture) or stationary phase (36h of culture) or after 72h growth (“aging coccoids”) was compared to PG extracted from the pA1-bearing strain either 8h or 24h after induction (toxin-induced coccoids, 24h and 36h total culture time) and from the pA1* strain as a negative control. Samples were digested with mutanolysin and subjected to HPLC/MS analyses. The relative abundance of muropeptides in each sample was calculated according to Glauner *et al*. (30) (**Table S1**). We found that the PG of B128 WT stationary phase and “aging coccoids” as well as the pA1 induced coccoids present enhanced amounts of GlcNAc-Mur*N*Ac-dipeptide (GM2) as compared to helical exponential phase *H. pylori* and to the negative control with pA1*. This GM2 increase seems to be at the expense of the GM3 muropeptides, whose amount is reduced in “aging coccoids” and in pA1-induced bacteria where it reaches a minimal value 24h post-induction. This shows that during the transition from exponentially growing spiral bacteria to coccoids, the muropeptide composition undergoes important modifications that are comparable for the toxin-induced coccoids and “aging coccoids”. These changes, compatible with a looser PG macromolecule, suggest that the coccoid formation under these two conditions might be generated by similar mechanisms.

### Kinetics of the toxin-induced *H. pylori* morphological transformation assessed by live cell imaging

To progress in our understanding of the morphological conversion of *H. pylori*, we monitored the entire process of *H. pylori* toxin-induced conversion by live cell time-lapse microscopy under physiological microaerobic conditions in a temperature-controlled chamber. The *H. pylori* B128 strain tested constitutively expressed cytoplasmic GFP, was impaired in motility (by deletion of *flaA* encoding the major flagellar protein) and carried the inducible toxin-expressing plasmid pA1. Individual live *H. pylori* bacteria were monitored in agarose pads by both bright field and fluorescence microscopy.

First, the general growth features of the strain between cell birth and its next division were established on 61 bacteria in the absence of the IPTG inducer. Under these conditions, statistics of the data collected showed that the strain has a mean doubling time of 165 min (indicative of optimal growth conditions) with an initial bacterial length of 1.9 µm after division and a maximum length prior to division of 3.2 µm (**Fig. S4**). In a second round of experiments, the A1 toxin was induced by IPTG addition to the pad and snapshots of the cells were taken at intervals of 10 min during 10h (**Fig. 4**). A total number of 158 cells were analyzed from 2 independent experiments that gave consistent results. Three representative movies can also be viewed in **Fig. S5**. During the 10h of analysis, about 70% of bacteria showed visible cell shape morphological transformations; these bacteria were further analyzed. To quantify the changes in shape of individual *H. pylori* cells over time, we used the MicrobeJ software (28) and applied circularity attributes to discriminate the populations. As illustrated in **Fig. 4**, a circularity value of 0.7 defines the first steps of *H. pylori* cells bending that is followed by quasi-round bacteria (circularity >0.8) and finally fully developed coccoids (circularity >0.9).

**Figure 4:**
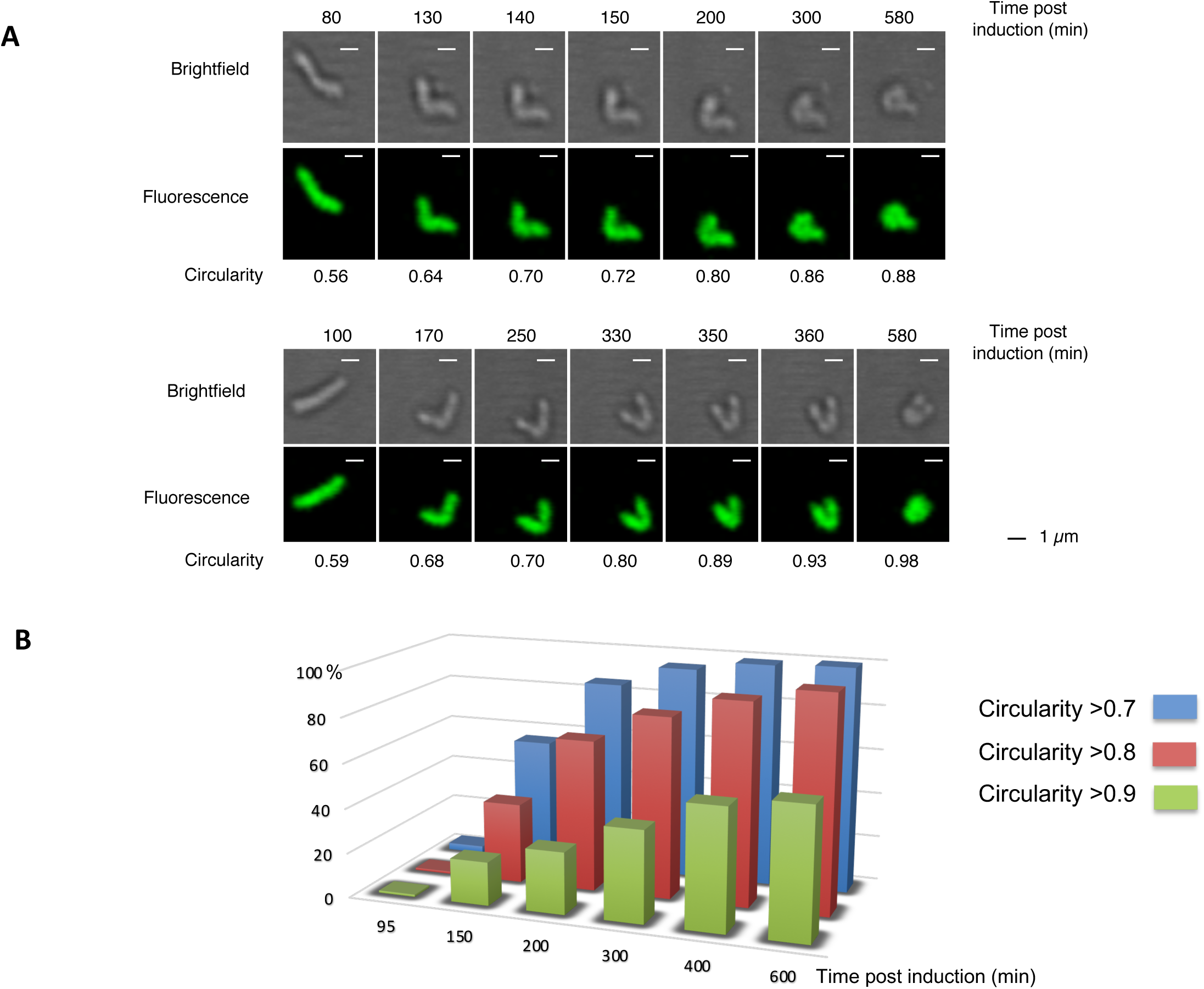
Time-lapse microscopy of *H. pylori* conversion to coccoid upon toxin expression. Live cell time lapse microscopy was used to record, during 10 hours every 10 min, the morphological modifications of individual *H. pylori* B128 strain cells after IPTG-induction of the AapA1 toxin. Representative movies of the transformation can be seen in **Fig. S5**. **A) Bright field and fluorescence images of two examples of these kinetics**. Circularity (>0.7; >0.8; >0.9) of the bacteria was measured with the MicrobeJ software. Scale bar represents 1µm. **B) Quantification of the evolution of the formation of round *H. pylori* cells over time after IPTG induction**. The percentage of each type of bacterium was classified as a function of its circularity (>0.7; >0.8; >0.9).

The changes in circularity were measured for 110 individual bacteria over time. **Fig. 4** provides an overall view of the progressive rounding of the analyzed bacteria. The kinetics of circularity changes reveal that the first step (>0.7) starts as early as at a median of 150 min post-induction, which is very close to the doubling time of *H. pylori* cells. Most interestingly, the length of bacteria just before they enter the first morphological transformation (circularity >0.7) was 3.46 µm (±0,6), which is very close to the maximum length we measured before *H. pylori* division. In addition, we observed many examples of two bacteria that were dividing but not yet separated and from which one out of the two underwent morphological transformation (see movies in **Fig. S5**). This analysis gave us a dynamic picture of the morphological transformation of *H. pylori*. Importantly, it suggests that the observed morphological transformation is likely to be consecutive to a toxin-induced perturbation of cell elongation and division, through a mechanism that is still to be defined.

### Toxin-antitoxin imbalanced expression upon oxidative stress

We then searched for the physiological signal that could trigger the induction of the AapA1 toxin expression in *H. pylori*. First, the respective activities of individual *aapA1* and *IsoA1* promoters were measured with chromosomally expressed transcriptional *lacZ* fusions (**Fig. S1**). Under normal growth conditions, ß-galactosidase activities indicated that the antitoxin promoter has a 10-fold stronger activity than the toxin promoter (**Fig. 5A**). This result is consistent with type I TA features where the antitoxin RNA is strongly expressed (but highly labile) compared to the more stable toxin transcript that is expressed at a lower level (8, 9, 22). The *lacZ* fusions were then used to follow the activity of the two promoters during *H. pylori* growth and to search for conditions relevant to *H. pylori* life-style that could lead to imbalanced expression of the two promoters. Only a slight increase in activity of both promoters was observed over time of growth (**Fig. 5A**). No significant differential expression of the promoters was observed upon acid, antibiotic stress or during exposure to high nickel concentrations (**Fig. S6**). In contrast, oxidative stress induced by hydrogen peroxide (H_2_O_2_) resulted in a strong specific decrease in the *IsoA1* promoter activity while, at the same concentration, the toxin promoter activity was marginally reduced (**Fig. 5B**). Exposure to paraquat, another oxidative stress agent, resulted in a comparable strong decrease of both promoters. The expression patterns and stability of the *aapA1* and *isoA1* transcripts were then analyzed by Northern blot with total RNA extracted at different time points after rifampicin addition. As shown in **Fig. 5C** and **Fig. S7**, the *aapA1* transcript was detected as a 175 nucleotide-long band with an estimated half-life of approx. 120min. In contrast, the 75 nt *isoA1* full length transcript declines much faster, with an estimated 25min half-life, and the production of an approx. 50 nt short processed form. Upon hydrogen peroxide exposure, the half-life of the AapA1 transcript was not significantly modified while we observed a more rapid depletion of the full-length *IsoA1* transcript with an almost immediate accumulation of the processed form (**Fig. 5C**). From three independent experiments, we quantified the relative amounts of the transcripts under normal growth conditions versus H_2_O_2_ exposure during 120 min after rifampicin addition (**Fig. 5D**). No significant change was observed for the *aapA1* transcript while H_2_O_2_ exposure resulted in a diminished half-life of the *IsoA1* full length transcript and in an imbalanced ratio of *IsoA1* in favor of its processed form (**Fig. 5D**). Taken together, these results show that H_2_O_2_ causes both diminished *IsoA1* transcription and increased degradation of *IsoA1* full-length transcript. These observations suggest that, by decreasing the amounts of antitoxin transcript, exposure to oxidative stress favors translation of the AapA1 mRNA and thus toxin production that ultimately leads to coccoid transition.

**Figure 5:**
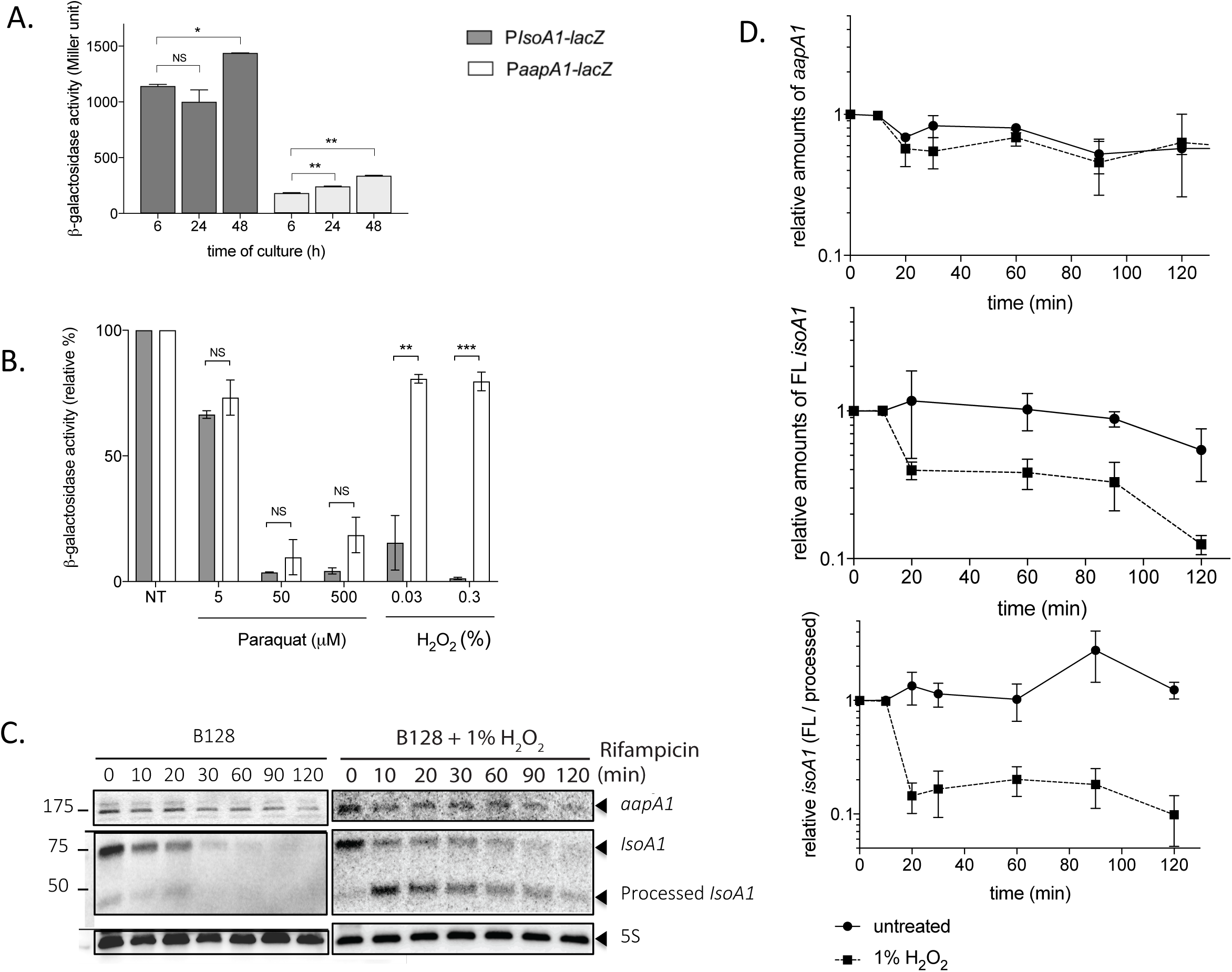
Oxidative stress generated by hydrogen peroxide results in decreased antitoxin promoter activity and promotes *IsoA1* transcript processing. **A) Activity of the AapA1 and IsoA1 promoters as a function of growth**. β-galactosidase activities (expressed in Miller units) measured with *H. pylori* B128 strain carrying chromosomal P*aapA1-lacZ* and P*IsoA1-lacZ* fusions after 6, 24 and 48h growth. Results from 3 independent experiments performed in duplicates are shown. Error bars represent the standard deviation, with * corresponding to *P* < 0.05, and ** to *P* < 0.01, indicating that the mean values are significantly different, and NS indicating that they are not significantly different (*P* >0.05). The activity of the toxin fusion (*aapA1-lacZ*) is about ten times that of the antitoxin fusion (*IsoA1-lacZ*) and the activity of both promoters slightly increases as a function of growth. **B) Oxidative stress generated by H**_**2**_**O**_**2**_ **decreases antitoxin promoter activity**. β-galactosidase activities expressed by P*aapA1-lacZ* and P*IsoA1-lacZ* fusions were measured after 6h treatment with oxidative stress generators, paraquat (5, 50 and 500 µM) and H_2_O_2_ (0.03 and 0.3%). β-galactosidase activities are presented as a ratio (expressed in %) of activities measured with stress versus activities of untreated samples. Hydrogen peroxide strongly decreases the expression of the P*IsoA1-lacZ* fusion. Results from 3 independent experiments performed in duplicates are shown. Error bars represent the standard deviation, with ** corresponding to *P* < 0.01 and *** to *P* <0.001, indicating that the mean values are significantly different, and NS indicating that they are not significantly different (*P* >0.05). **C) Determination of AapA1 mRNA and IsoA1 RNA half-lives in *H. pylori* strain B128 under normal conditions and upon hydrogen peroxide exposure**. Northern blots of total RNA from B128 WT strain grown under normal conditions or exposed to 1% hydrogen peroxide, extracted at the indicated times after addition of 80 μg/ml rifampicin. Five µg of RNA were loaded in each lane and the membranes were probed with [γ-32P] ATP-labeled oligonucleotides specific to the following RNAs, *aapA1, IsoA1* and 5S rRNA as a loading control. In the presence of H_2_O_2_, the half-life of *IsoA1* transcript is diminished in favor of the processed *IsoA1* form. **D) Hydrogen peroxide promotes *IsoA1* transcript processing**. Graphic representation of the effects of hydrogen peroxide exposure on the amounts of *aapA1* and *IsoA1* transcripts as well as on *IsoA1* processing during 120 min after rifampicin addition. The relative amounts of *aapA1* (upper graph) and full length (FL) *IsoA1* (middle graph) RNAs versus 5S rRNA are shown. The lower graph presents the relative amounts of the two forms of *IsoA1*, full length (FL) versus the processed form. The results of 3 independent experiments are shown, error bars represent the standard deviation analyzed by two-way ANOVA multiple comparisons.

### Oxidative stress induction of coccoids: are class A Type I TA the only effectors?

Since we established that oxidative stress results in imbalanced expression in favor of the expression of the AapA1 toxin, we examined the consequences of exposure to H_2_O_2_ on *H. pylori* morphology and viability (**Fig. S8**). Upon treatment with 2% H_2_O_2_ of WT B128 strain, we measured a partial conversion to coccoids (around 25%) either after 6h or 24h. Treatment with 5% H_2_O_2_ resulted in a rapid and massive transformation (about 90% of cells) of this strain into coccoids after 6 or 24h without detectable cell lysis (**Fig. S8A**). Strain B128 contains six class A type I TA systems (**Fig. S9**). To assess their role, we constructed mutants carrying deletions of each of its active AapA/IsoA TA loci, *ΔA1, ΔA3, ΔA4* (including deletion of both *A4-1* and *A4-2* tandem systems), *ΔA5, ΔA6* or of the five loci collectively (*Δ5*) using a non-marked deletion strategy (31). A locus corresponding to the position of the A2 locus in other *H. pylori* strains was identified but its corresponding A2 ORF was inactive in strain B128. Under normal conditions, the growth rate, culturability and kinetics of occurrence of coccoids of the different mutants were comparable to that of the parental strain (**Fig. S8**). Most interestingly, treatment of the Δ5 mutant with 2% H_2_O_2_ revealed minimal conversion after 6h of treatment with only 10% of the cells identified as coccoids as compared to 25% coccoid conversion for the wild type B128 strain. After 24h treatment, about 40% of the Δ5 cells were coccoids. Upon treatment with 5% H_2_O_2_, both WT and Δ5 mutant strain displayed similar kinetics of conversion to coccoids (**Fig. S8A**). This suggests that, in the Δ5 strain, there is a significant delay in oxidative stress-induced coccoid formation.

We conclude that oxidative stress is triggering transformation into coccoids through type I class A TA systems but that other effectors are able to eventually drive this conversion. In some cases TA systems have been shown to play a role in persister cell formation under stress conditions (8, 10). Therefore, the number of persisters after exposure to hydrogen peroxide was measured for the ΔA1 and Δ5 mutants and compared to the parental strain (**Fig. S8**). No significant difference was observed, suggesting that, upon hydrogen peroxide stress, the class A TA systems are not promoting viable persisters in *H. pylori*.

## Discussion

Here we established that the expression of a small toxin from a toxin-antitoxin system triggers rapid morphological transformation of the spiral-shaped *H. pylori* bacterium into round coccoid cells. Transformation of spirals into coccoids has been observed after extended *H. pylori* growth (>70h, “aging coccoids”) or upon stress conditions (17) but the process itself was poorly characterized. Here we found that the morphological transformation starts as soon as 2.5h after toxin induction and that around 50% and 70% of the bacterial population convert into round coccoids depending on the conditions (liquid culture vs agar pads for the live imaging). In liquid culture, the amount of coccoids is probably underestimated since the vast majority of the cells had lost culturability suggesting that they are indeed entering the process of dormancy. We consider that the amount of 70% coccoid conversion that we observe by live microscopy is a more accurate value since in these experiments the shape evolution of single cells was precisely followed.

The toxin (AapA1) is a small hydrophobic peptide of a type I TA system that we showed here to target *H. pylori* inner membrane without being secreted. Toxins of type I TA systems from other organisms such as *Escherichia coli, Bacillus subtilis* or *Staphylococcus aureus* have been shown to localize to the inner membrane but there are no reports on induction of major cell shape modifications (4, 21, 32, 33). The *E. coli* TisB (34) and the *S. aureus* PepA1 and PepA2 (35, 36) toxins act by creating membrane pores (similar to phage holins) thereby disrupting the membrane potential, drastically impairing ATP synthesis and depending on the toxin concentration, either leading to the formation of persisters or to cell death (9, 33). One notable exception is the BsrG/SR4 Type I TA system of *B. subtilis* (32). The BsrG toxin was found to target the membrane without causing destruction or affecting the Proton Motive Force (PMF) but it rather directly interferes with the cell envelope biosynthesis, indirectly delocalizes the cell wall synthesis machinery and ultimately triggers bacterial autolysis (32). There are now two examples of type I TA hydrophobic toxins (BsrG and AapA1 reported here) that do not act by permeabilizing the membrane and probably reflect a novel mode of action for small peptidic toxins.

Using cryo-EM on *H. pylori*, we observed that toxin-induced *H. pylori* coccoids present neither visible membrane disruption nor pores even 8h post toxin induction and are ultrastructurally comparable to 70h-old “aging coccoids”. In addition, the membrane potential of coccoids was not dissipated. This is in contrast with a previous study that reported a total loss of membrane potential in *H. pylori* “aging coccoids”, associated with a complete loss of membrane integrity (37). Unlike the effects of the TisB toxin in *E. coli* (34), we found that the ATP content in our *H. pylori* toxin-induced coccoids is marginally affected and drops only after prolonged culture, as is the case for spiral-shaped cells (38).

Toxins expressed by TA systems cause growth arrest by interfering with conserved vital cellular processes such as translation, division or PG synthesis (9, 33). In bacteria, PG cell wall dictates cell shape. We found that coccoid transformation was accompanied by changes in PG composition. Indeed, when compared to spirals, coccoids have a three-fold increase in dipeptide monomers concomitant with a tripeptide monomers reduction. These changes are comparable to those previously reported for “aging” *H. pylori* coccoids (39) and were confirmed here. Our results point to a reduction in the potential for generating new cross-bridges in the PG; this is compatible with a looser PG macromolecule and could explain the morphological transition to coccoids.

Live imaging analysis of the toxin-induced conversion of *H. pylori* spirals into coccoids suggested that the AapA1 toxin could interfere with cell elongation and division. In *E. coli*, the CbtA toxin of a type IV TA system was found to inhibit cell division and cell elongation via direct and independent interactions with FtsZ and MreB (40). More work is needed to identify the molecular target(s) of the AapA1 toxin.

Under normal conditions, as for all TA systems, the *IsoA1* antitoxin potently inhibits the AapA1 toxin, by preventing its expression through a multilayered mechanism (24, 25). We searched for physiological conditions that could result in toxin expression. We found that oxidative stress, generated by H_2_O_2_, causes a rapid and specific decline in the levels of IsoA1 antitoxin full-length transcript by reducing both its promoter activity and enhancing its degradation through a mechanism that remains to be precisely defined. Imbalanced expression of the AapA1/IsoA1 system in favor of the toxin mRNA should lead to toxin expression. Accordingly, we observed that H_2_O_2_ causes a rapid conversion of *H. pylori* into coccoids resembling those induced by the AapA1 toxin. Prolonged exposure to aerobic conditions was previously reported to lead to coccoid formation, however, in that case these cells lost their membrane integrity (41). Regulation of type I TA systems in response to stress has been observed in other species. For the *bsrE*/SR5 type I system of *B. subtilis*, the SR5 antitoxin is affected by pH, anoxia and iron limitation while the BsrE toxin is sensitive to temperature shock and alkaline stress (42). In *S. aureus*, the SprA1/SprA1_AS_ system is induced in response to acidic or oxidative stresses (35) and the antitoxin RNAs of the SprG/SprF systems are reduced upon oxidative stress exposure (43). In *E. coli*, several type I toxins are induced by the SOS response (8), a system that does not exist in *H. pylori*. The biological function of *H. pylori* coccoids is still a matter of debate. Coccoids are observed in prolonged *in vitro* cultures and induced by stress conditions, mostly aerobic, anaerobic culture or exposure to antibiotic or oxidative stress. However, these reports are difficult to compare since no standard procedure was used and analysis was most of the time performed after more than a week exposure to stress (17, 37, 44). *H. pylori* coccoids are non-culturable cells under standard laboratory growth conditions. Our analysis of both toxin-induced and 70h “aging coccoids” are in favor of their viability despite their “unculturability”. We argue that contradictory conclusions on viability refer to “damaged” or so-called “fragmented” coccoid forms corresponding to membrane-less bacteria in prolonged 5 days-old *H. pylori* cultures (37). In favor of coccoid viability are previous reports showing that coccoids express virulence factors (17) and are capable, like spirals, of binding to host cells and inducing cellular changes (45). *H. pylori* coccoids were also visualized in human gastric biopsies (46), probably as part of biofilms (44); however it cannot be excluded that they correspond to a transverse view of a spiral bacterium. In mice, coccoids induce gastritis (47) and can revert to colonizing spiral bacteria (48). Taken together, these observations suggest that coccoids are “dormant” viable forms of *H. pylori* that recover during host infection and might play a role in transmission or in treatment failure. In agreement with this view, Chaput *et al*. (49) showed that coccoids present decreased activation of NF-κB and might allow the bacterium to escape the host immune response.

We showed that *H. pylori* coccoids are induced by a class A type I TA toxin and by oxidative stress probably through the imbalanced expression of TA systems. During infection, *H. pylori* is indeed exposed to harsh oxidative stress as a consequence of the chronic inflammation it generates (50). Deletion of the five “class A” Type I TA clusters of *H. pylori* was found to delay the oxidative stress induction of coccoids but did not preclude it nor did it change the number of persister cells. We concluded that these toxins are probably not the only triggers of coccoid transformation and suggest that the two other classes of chromosomally-encoded type I TA systems (B and C representing 9 systems) might also be important.

Dormant/persister bacteria present enhanced tolerance to antibiotics and in some cases, have been shown to be induced by TA systems (8, 10), although other mechanisms resulting in lowered ATP content and/or lowered PMF are also associated with their occurrence (13, 14). We hypothesize that TA-induced *H. pylori* coccoids are dormant bacteria as defined in a recent consensus review (51). As previously proposed, non-culturable cells and “classical” dormant bacteria are part of a shared “dormancy continuum” (52). In our model, this continuum would depend on the intracellular toxin concentration as already suggested (53).

Stress-induced morphological transformation of bacteria into non-culturable coccoid-like cells has been reported in at least 85 bacterial species among which are important bacterial pathogens such as *Campylobacter jejuni, Vibrio cholerae* or *Salmonella typhimurium* (19). For these organisms, the inducing trigger and function remains to be characterized. We postulate that these forms are dormant bacteria and that our findings on *H. pylori* might be more generally relevant and play important roles in recurrent and persistent bacterial infections.

## Acknowledgments

This work was funded by the Agence National de la Recherche (ANR 09 BLAN 0287 01 and ANR 12 BSV5-0025-02), the Laboratoire d’Excellence IBEID (Integrative Biology of Emerging Infectious Diseases) Grant ANR-10-LABX-62-IBEID from the French government’s Investissement d’Avenir program and the Pasteur-Weizmann Consortium of “The Roles of Noncoding RNAs in Regulation of Microbial Life Styles and Virulence”. LEM was funded by a Roux fellowship of the Institut Pasteur and by a Carnot MI fellowship. ATA is part of the Pasteur Paris University (PPU) International PhD Program. This project has received funding from the European Union Horizon 2020 research and innovation program under the Marie Sklodowska-Curie grant agreement No 665807, and from the Institut Carnot Pasteur Microbes and Santé. AR was supported by post-doctoral fellowship from the Labex IBEID (10-LABX-62-IBEID). We also thank Janssen for financial support. We are grateful to F. Darfeuille, I. Iost and H. Arnion for the gift of plasmids and to W. Fischer for the gift of anti-AlpA antibodies. We also appreciate the expertise and help of A. Chery-Faleme, M. Fromont-Racine and A. Jacquier with Northern blotting, of R. Wheeler for peptidoglycan extraction, of Z. Baharoglu and I. Santecchia for flow cytometry and of A. Ducret for image analysis. We thank M. Denic and S. Kumar for comments on the manuscript and J. Berry for discussions. Finally, we thank M. Nilges and the Equipex CACSICE for providing the Falcon II direct detector.

## Material and Methods

### Bacterial strains and growth conditions

The *H. pylori* strains used in this study (Suppl **Table S2**) were all derived of strain B128 (54, 55). Plasmids (suppl **Table S3**) used to create or complement *H. pylori* mutants were constructed and amplified using *E. coli* One-Shot TOP10 or DH5α strains (Thermofisher). *H. pylori* strains were grown on Blood Agar Base 2 (Oxoid) plates supplemented with 10% defibrinated horse blood and with the following antibiotics-antifungal cocktail: amphotericin B 2.5 μg.ml^-1^, polymyxin B 0.31 μg.ml^-1^, trimethoprim 6.25 μg.ml^-1^ and vancomycin 12.5 μg.ml^-1^. Selection of *H. pylori* mutants was performed using kanamycin 20 μg.ml^-1^, Streptomycin 10μg.ml^-1^, Apramycin 10μg.ml^-1^ or chloramphenicol 8 μg.ml^-1^. For liquid cultures, we used Brain Heart Infusion (BHI) broth (Oxoïd) supplemented with 10% Fetal Calf Serum (FCS) (Eurobio), the antibiotics-antifungal cocktail and the selective antibiotic when necessary. *H. pylori* cells were grown at 37°C under microaerophilic atmosphere conditions (6% O_2_, 10% CO_2_, 84% N_2_) using an Anoxomat (MART Microbiology) atmosphere generator. When indicated, 1mM of isopropyl ß-D-1-thiogalactopyranoside (IPTG, EuroMedex) was added to agarose pads or culture media.

### Molecular techniques

Molecular biology experiments were performed according to standard procedures and the supplier (Fermentas) recommendations. NucleoBond Xtra Midi Kit (Macherey-Nagel) and QIAamp DNA Mini Kit (Qiagen) were used for plasmid preparations and *H. pylori* genomic DNA extractions, respectively. PCR were performed either with Taq Core DNA polymerase (MP Biomedicals), or with Phusion Hot Start DNA polymerase (Finnzymes) when the product required high fidelity polymerase. The pGEM-T easy vector systems (Promega) was used to construct, in *E. coli*, the suicide plasmids that served for markerless deletions of TA systems in *H. pylori*.

### Construction of H. pylori strains carrying mutations or plasmids

Mutations of *H. pylori* were introduced into strain B128 either WT or a streptomycin resistant variant B128 *rpsL1* for markerless mutagenesis (31, 56). Chromosomal deletions of the *AapA1/*IsoA1 locus, of the *flaA* gene, or of the different class A TA loci were performed either by insertion of a selectable antibiotic resistance marker to disrupt or replace the gene of interest or when necessary, by the markerless counter-selected mutagenesis strategy (31, 56). Plasmids were derived from the pILL2157 *E. coli/H. pylori* shuttle vector that contains an IPTG-inducible promoter (27). Deletions were introduced by allelic exchange using suicide plasmids (see suppl **Table S3**) or PCR fragments. Introduction of plasmids and construction of *H. pylori* mutants were obtained by natural transformation and selection with the corresponding antibiotic as described previously (31). PCR and sequencing of the regions of interest were used to validate the introduction of plasmids, deletion of genes of interest and correct insertion of fusions. Primers used for these constructs or their validation are listed in Suppl **Table S4**.

### Fluorescence Microscopy

Fluorescence microscopy was performed with an Axio Observer microscope (Zeiss) equipped with an Axiocam camera under an X100 magnification. Acquisition images was performed using the axiovision software. Images were cropped and adjusted using ImageJ v2.0.0 software. Cells were quantified and assigned to helical or coccoid categories based on phase contrast images by using the MicrobeJ plugin of ImageJ (28) with the following parameters: (i) helical cells had an area of more than 0.7 μm^2^, a length between 1 and 5 μm, a width between 0.4 and 1 μm and a circularity of less than 0.8, and (ii) coccoid cells had an area between 1 and 2 μm^2^, a width between 0.4 and 1.7 μm and a circularity greater than 0.7.

*H. pylori* B128 strains were grown in BHI medium with chloramphenicol (8 µg/ml) to maintain the plasmids. 1 mM IPTG was added to the culture and samples were taken at different time-points, concentrated by 2 min centrifugation at 3000 x g and washed twice with PBS buffer. Cells were immobilized using 2% (wt/vol) agarose pads containing PBS before being imaged. In **Fig. 1C**, cell membranes were stained with 4% FM4-64 (Thermofisher Scientific). In **Fig. 2B**, cell membranes were stained with 0.01 mM TMA-DPH (1-(4-trimethylammoniumphenyl)-6-phenyl-1,3,5-hexatriene p-toluenesulfonate, Euromedex).

### Flow cytometry for proton motive force (PMF) measurement

*H. pylori* strains were grown in triplicate for 16h and then treated or not with 1 mM IPTG. Samples were taken at 0, 8 and 24h post-induction and 10^7^ live cells per experimental condition were washed and stained with 25 nM MitoTracker® Red CMXRos (Invitrogen), a PMF-sensitive dye. As a control for PMF depletion, pA1* cells were treated with 500µM of the protonophore TCS (3,3’,4’,5-Tetrachlorosalicylanilide; Fisher scientific). The fluorescent signal from at least 100,000 individual bacteria per condition was measured by flow cytometry with a MCSQuant® VYB analyzer (Miltenyi Biotec) (Y2 channel, λex=561 nm and λem=605-625 nm) after calibration. The experiment was performed three times. Data were analyzed with FlowJo V10.

### Fractionation and Immunoblotting

The protocol of *H. pylori* fractionation was adapted from (57). *H. pylori* strains were grown in the absence or presence of 1mM IPTG. When cultures reached OD_600_ 0.8, cells were harvested by centrifugation, washed twice in phosphate-buffered saline medium (PBS) prior to their disruption by sonication in Buffer 1, 10mM Tris-HCl, Ph 7.5 containing 5mM ß-mercaptoethanol and proteases inhibitors (cOmplete^™^, EDTA-free Protease Inhibitor Cocktail, Roche). Cell debris were removed by centrifugation (10 min, 16,000 x *g*, 4°C) and supernatants containing the soluble extract and membrane fractions were collected as total extracts. Samples of total extracts were frozen and the remaining supernatants were subjected to ultracentrifugation (125,000 x *g*, 45 min, 4°C). Samples from the supernatant containing the cytoplasm and periplasm were collected and frozen while pellets containing total membranes were resuspended in buffer 1 supplemented with 1% N-lauroylsarcosine (buffer 2) prior to being ultracentrifuged (125,000 x *g*, 45 min, 4°C). Samples of inner membrane fractions from supernatants were frozen. Outer membrane pellets were washed twice in buffer 2 to avoid inner membrane fractions contamination prior to freezing. For each cellular compartment, equal cellular amounts were loaded and separated on a 4-20% Mini-Protean TGX precast protein gel (BioRad) and subsequently electrotransferred on a polyvinylidene difluoride (PVDF) membrane (Biorad). The control *H. pylori* proteins PPB2, AlpA and AmiE and NikR as well as the fusion protein A1-GFP were detected with rabbit polyclonal antibodies α-PBP2 (58), α-AlpA (57), α-AmiE (59) and α-NikR (60); with goat anti-GFP-HRP antibody (Abcam) at the respective dilutions of 1:2,000, 1:1,000, 1:100, 1:5,000 and 1:5,000. Goat anti-rabbit IgG-HRP (Santa Cruz) was used as secondary antibody (1:10,000 dilution) and the detection was achieved with the ECL reagent (ThermoFisher Scientific).

### ATP extraction and assay

Exponentially growing *H. pylori* cells (6h of culture) and stationary growing cells (16h) were harvested by centrifugation at room temperature for 4 min at 5000*g*. Metabolites from the resulting cell pellets were extracted immediately using 300μL of a solvent mixture of Acetonitrile/Methanol/H_2_O (40/40/20) for 15min at 4°C and 10min at 95°C. Mixtures were subsequently spun in a microfuge for 5 min at maximum speed and 4°C to separate insoluble materials from the extracted metabolites. The resulting pellets were then re-extracted twice with 200 μL of solvent at 4°C. The supernatants were pooled to yield 700 μL of final extract. Metabolites were lyophilized and subsequently diluted in water for ATP assays. ATP content was determined by a luciferase-based ATP bioluminescence assay kit (BacTiter-Glo™ Microbial cell viability assay, Promega). Luminescence values were determined using a 10 sec RLU signal integration time and measured using a Centro XS^3^ LB960 Luminometer (Berthold Technologies). ATP concentrations were calculated based on values determined using serial dilutions of known amounts of ATP and expressed as a function of the OD_600nm_ of the corresponding culture.

### Cryo-electron microscopy

Four µL of *H. pylori* bacteria were spotted on glow-discharged lacey grids (S166-3, EMS) or Quantifoil R2/2 (Quantifoil, Germany). The samples were cryo-fixed by plunge freezing at −180°C in liquid ethane using a Leica EMGP (Leica, Austria). Grids were observed at 200kV with a Tecnai F20 (Thermo Fisher Scientific). Images were acquired under low-dose conditions using the software EPU (Thermo Fisher Scientific) and a direct detector Falcon II (Thermo Fisher Scientific).

### Time-lapse microscopy

Confocal analysis of live cells was performed as previously described (61). Twenty µL of exponentially grown bacteria (10^5^ cells) suspended in BHI with or without 1mM IPTG were deposited on 35mm glass bottom Petri dishes. The suspension was covered with BHI medium in 1.5% low melting agarose supplemented with Chloramphenicol (8µg/µl) with or without IPTG (1mM). Live-cell imaging was performed with a Nikon A1R confocal laser scanning microscope system attached to an inverted ECLIPSE Ti (Nikon Corp., Tokyo, Japan) and equipped with an environmental chamber allowing the control of temperature (37°C), humidity (90%) and gas mixture (10% CO_2_, 3% O_2_). GFP fluorescence images were captured through a Plan APO 60X objective (NA: 1.40) by using optimal spatial resolution settings. The cytoplasm compartment volume was defined by using GFP staining (excitation with 488 nm laser, emission collected with an 500/50 nm filter set). Image captions were performed every 10min during 10h. Image treatment and analysis were performed using NIS elements (Nikon Corp., Tokyo, Japan), and ImageJ and MicrobeJ software (28).

### ß-galactosidase activity assays

B128*ΔAapA1-IsoA1::PaapA1-lacZ-Kan* and B128*ΔAapA1-IsoA1::PIsoA1-lacZ-Kan* strains were grown to OD_600nm_ 0.3 in BHI liquid medium, then divided into two samples, one of them being submitted to stress conditions during 6h with 5, 50 and 500 µM paraquat (Sigma), 0.03 and 0.3% hydrogen peroxide (Sigma), 20 and 200 mM NiCl_2_ (Sigma), acidic pH, antibiotics with 0.1 and 1 mg/mL tetracycline, or 0.05 and 0.5 mg/ml rifampicin. Then 0.5ml samples were taken, washed twice with 1X PBS (Phosphate Buffered Saline) and further permeabilized with 100µL Chloroform and 50µl SDS 0.1% in Z buffer containing 70mM Na_2_HPO_4_, 30mM NaH_2_PO_4_, 1mM MgSO_4_ and 0.2mM MnSO_4_ (62). Samples were briefly vortexed and incubated at 28°C for 2 min. The ß-galactosidase assay was started by adding 0.1mL ONPG (ortho-nitrophenyl-β-galactoside, ThermoFischer) at 4mg/ml and stopped by the addition of 0.5mL 1M Na_2_CO_3_ when sufficient yellow color was reached. The ß-galactosidase activity is expressed in Miller units (62) and represented in **Fig. 5** and **S6** as percentage of activities relative to the control activity with the corresponding strains not exposed to stress and was calculated from 3 independent experiments performed in duplicates.

### Total RNA extraction and Northern blotting

Total RNAs were extracted from 10 mL *H. pylori* cultures (OD_600_ 0.5 −0.9) exposed or not to 1% H_2_O_2_ using the NucleoSpin miRNA kit (Macherey Nagel) at different time points after 80 µg/mL rifampicin addition (to stop transcription). For Northern blotting, 5 µg of total RNA were separated on 10% Mini protean TBE Urea gel (Biorad) and transferred to Hybond N+ (Amersham Biosciences) membrane using a Trans-Blot Turbo system (Biorad). A denaturated DNA marker was used for size estimation.

Transferred RNA was fixed to the membrane by UV irradiation for 2 min. The membrane was blocked for 45 min at 42°C with ULTRAhyb Hybridization Buffer (Ambion), then 20 µl of 5’-labeled (γ^32^P) oligodeoxynucleotides (**Table S4**) were added and the membrane was further incubated overnight at the same temperature. After three washes for 10 min at 65°C with 2x SSC 0.2 % SDS, the membrane was exposed to a phosphorimager screen (KODAK) and scanned with FLA-9000 Phospho Imager (Fujifilm).

### Peptidoglycan extraction and Muropeptides analysis by HPLC/HRMS (high resolution mass spectrometry)

Samples corresponding to 2 mL of culture at OD_600nm_ 10 of *H. pylori* were taken and chilled in an ice-ethanol bath, each condition was prepared in triplicates. The samples were B128 WT strain in exponential phase (24h culture); stationary phase (36h culture); after 72h culture (“aging coccoids”). To examine the effect of the A1 toxin, we performed a culture of 16 h, added IPTG and then took samples of the following strains: B128 ΔaapA1-IsoA1 + pA1 strain after 8h of induction (equivalent to 24h of total culture time) or 24 h of induction (equivalent to 36h); and as a negative control, of strain B128 ΔaapA1-IsoA1 + pA1* after 8h and 24 h of induction (total 24h and 36h, respectively). Crude murein sacculi were immediately extracted in boiling sodium dodecyl sulfate (SDS 4% final) (63). The resulting purified peptidoglycan was digested overnight at 37°C in 12.5 mM sodium phosphate buffer (pH 5.6) supplemented with 100 UI of mutanolysin from *Streptomyces globisporus* ATCC 21553 (Sigma). The reaction was stopped by heat inactivation of the enzyme and insoluble material was removed by centrifugation. Soluble muropeptides were then reduced with sodium borohydride in borate buffer (pH 9). After centrifugation, reduced muropeptides were diluted 20-fold in water-formic acid 0.1% (v/v; solvent A). HPLC/HRMS (High Performance Liquid Chromatography/High Resolution Mass Spectrometry) was performed on an Ultimate 3000 UHPLC (Ultra High-Performance Liquid Chromatography) system coupled to a quadrupole-Orbitrap mass spectrometer (Q-Exactive Focus, Thermo Fisher Scientific). Muropeptides were separated on an Hypersil Gold aQ analytical column (1.9 µm, 2.1×150 mm) at 200 µL/min, column temperature at 50°C. Applied linear gradient from 0 to 12.5% acetonitrile + 0,1% (v/v) formic acid (solvent B), followed by increasing to 20% B at 25 min for 5 min, hold 20% B for min and additional 10 min with 100% A for column re-equilibration. Eluted muropeptides were introduced into Q-Exactive instrument, operating in positive ion mode, and then were analyzed in the data-dependent acquisition mode (FullMSddMS2). The identification of muropeptides was performed after precursor ion fragmentation (MS2 analysis). Data were then processed with the software TraceFinder 3.3 (Thermo Fisher Scientific) for peak areas determination, the relative abundance of muropeptides in each sample was calculated according to Glauner *et al*. (30).

### Statistical analysis

The Student’s t test was used to determine significance of the means of the data for all the experiments except for the Northern blots. Error bars represent the standard deviation, with * (*P* < 0.05), ** (*P* < 0.01), *** (P <0.001), **** (*P* < 0.0001) indicating that the mean values are significantly different and NS that they are not significantly different (*P* >0.05). SEM corresponds to standard error of the mean. The two-way ANOVA multiple comparisons test was used to compare Northern blots bands intensities under different conditions.

