## Supplementary Table S1-S2-S3-S4 for "A peptide of a type I toxin-antitoxin system induces *Helicobacter pylori* morphological transformation from spiral-shape to coccoids"

**Table S1: Summary of the muropeptide composition of peptidoglycan extracted from B128 WT strain during exponential phase (24h), early stationary (36h) and late stationary phase (72h culture) and from the B128 ∆*aapA1-IsoA1* + pA1 strain, 8 and 24 hours after toxin induction by IPTG addition and, as a negative control, from B128 ∆*aapA1-IsoA1* + pA1* 8h and 24 h after IPTG addition.**

**Panel A:** Summary of the muropeptide composition of peptidoglycan extracted from *H. pylori* B128 WT strain during exponential phase (24h, first column); stationary phase (36h, second column); after 72h culture ("aging" coccoids, third column); of toxin-induced coccoids of B128 ∆*aapA1-IsoA1* + pA1 strain after 8h of induction (equivalent to 24h of culture, fourth column) or 24 h of induction (equivalent to 36h of culture, fifth column); and as a control, of strain B128 ∆*aapA1-IsoA1* + pA1* after 8h of induction (sixth column) and 24 h of induction (seventh column). Each condition was analyzed in triplicates. The relative abundance of muropeptides in each sample was calculated according to Glauner *et al.* (30). Arrows show major statistically significant changes measured in the muropeptide composition when comparing exponential phase grown bacteria with aging coccoids and with A1 toxin induced coccoids (cultures at equivalent timepoints). These data show that both "aging coccoids" and toxin-induced coccoids present similar changes namely significant GM2 increase and GM3 reduction, which for the toxin-induced condition is already visible at 8h post-induction and is accentuated at 24h post-induction. In contrast, for the control condition, pA1*, no such changes are measured.

| | | Area - % for each muropeptide[a] | | | | | | |
|---|---|---|---|---|---|---|---|---|
| | | B128 | | | B128 Δ*aapA1-IsoA1* + pA1 | | B128 Δ*aapA1-IsoA1* + pA1* | |
| | | Exponential phase (24h) | Stationary phase (36h) | "Aging" coccoids (72h) | 8h post-induction (24h culture) | 24h post-induction (36h culture) | 8h post-induction (24h culture) | 24h post-induction (36h culture) |
| *Peak n°* | *Monomers* | 69.11 ± 1.00 | 73.11 ± 1.17 | 73.38 ± 0.23 | 59.83 ± 0.41 | 61.23 ± 0.58 | 58.75 ± 1.07 | 59.98 ± 1.04 |
| 3 | GM2 | 9.04 ± 0.57 | 24.67 ± 1.24 | 27.27 ± 0.54 | 15.62 ± 1.24 | 23.72 ± 1.21 | 6.55 ± 0.7 | 13.19 ± 3.51 |
| | | | ↗ | ↗ | ↗ | ↗ | | |
| 1 | GM3 | 9.42 ± 0.94 | 7.49 ± 0.71 | 4.67 ± 2.26 | 5.00 ± 1.3 | 1.12 ± 0.19 | 8.14 ± 0.72 | 5.51 ± 1.23 |
| | | | | ↘ | ↘ | ↘ | | |
| 4 | GM4 | 16.91 ± 0.34 | 10.8 ± 0.48 | 14.52 ± 2.77 | 9.43 ± 0.76 | 9.02 ± 0.23 | 11.97 ± 0.77 | 10.33 ± 1.39 |
| 5 | GM5 | 28.06 ± 0.8 | 23.92 ± 0.69 | 20.53 ± 0.54 | 25.99 ± 1.95 | 23.30 ± 0.6 | 28.16 ± 2.09 | 26.69 ± 0.23 |
| 2 | GM4+gly[b] | 5.69 ± 0.26 | 6.24 ± 0.44 | 6.41 ± 0.74 | 3.79 ± 0.35 | 4.07 ± 0.19 | 3.94 ± 0.13 | 4.25 ± 0.27 |
| | *Dimers* | 16.13 ± 0.2 | 13.63 ± 0.04 | 11.66 ± 1.32 | 15.41 ± 1.07 | 13.11 ± 0.71 | 15.40 ± 0.24 | 14.7 ± 1.94 |
| 7 | GM3 + GM4 | 2.21 ± 0.16 | 1.66 ± 0.3 | 1.41 | 2.43 ± 0.08 | 1.96 ± 0.11 | 2.52 ± 0.24 | 1.94 ± 0.41 |
| 8 | GM4 + GM4+gly | 0.93 ± 0.08 | 1.02 ± 0.01 | 1.01 ± 0.13 | 0.77 ± 0.04 | 0.69 ± 0.07 | 0.77 ± 0.06 | 0.81 ± 0.02 |
| 9 | GM4 + GM4 | 7.1 ± 0.19 | 5.84 ± 0.09 | 5.66 ± 0.76 | 5.73 ± 0.12 | 5.23 ± 0.17 | 6.11 ± 0.52 | 5.94 ± 0.15 |
| 10 | GM5 + GM4 | 5.89 ± 0.13 | 5.12 ± 0.37 | 4.29 ± 0.3 | 6.49 ± 0.97 | 5.22 ± 0.55 | 6.00 ± 0.38 | 6.01 ± 0.16 |
| | *Trimers* | | | | | | | |
| | GM4 + GM3 + GM4 | 0.06 ± 0.00 | 0.06 ± 0.01 | 0.12 ± 0.1 | 0.1 ± 0.01 | 0.09 ± 0.00 | 0.11 ± 0.01 | 0.10 ± 0.01 |
| | *Anhydromuropeptides* | 14.28 ± 0.88 | 12.8 ± 1.09 | 14.42 ± 1.19 | 22.63 ± 0.57 | 22.46 ± 0.18 | 23.77 ± 0.94 | 23.51 ± 0.63 |
| 12 | G(anhM)2 | 0.23 ± 0.02 | 0.54 ± 0.01 | 0.88 ± 0.01 | 0.7 ± 0.03 | 1.09 ± 0.06 | 0.65 ± 0.04 | 0.97 ± 0.29 |
| 6 | G(anhM)3 | 0.62 ± 0.04 | 0.54 ± 0.08 | 0.62 ± 0.16 | 1.01 ± 0.07 | 0.46 ± 0.03 | 1.06 ± 0.08 | 0.94 ± 0.1 |
| 11 | G(anhM)4 | 1.56 ± 0.13 | 0.84 ± 0.13 | 1.61 ± 0.07 | 1.59 ± 0.17 | 1.06 ± 0.02 | 2.29 ± 0.05 | 1.77 ± 0.46 |
| 13 | G(anhM)5 | 2.29 ± 0.2 | 2.06 ± 0.12 | 1.89 ± 0.37 | 2.49 ± 0.22 | 1.9 ± 0.04 | 3.00 ± 0.25 | 2.87 ± 0.11 |

| | | | | | | | | |
|---|---|---|---|---|---|---|---|---|
| 14 | G(anhM)4 + GM3 | 1.23 ± 0.11 | 1.06 ± 0.21 | 1.11 ± 0.1 | 2.52 ± 0.25 | 2.66 ± 0.15 | 2.38 ± 0.2 | 2.21 ± 0.3 |
| 15 | G(anhM)4+GM4+gly | 0.29 ± 0.03 | 0.41 ± 0.01 | 0.47 ± 0.01 | 0.48 ± 0.03 | 0.55 ± 0.02 | 0.44 ± 0.03 | 0.52 ± 0.2 |
| 16 | G(anhM)4 + GM4 | 3.63 ± 0.22 | 3.24 ± 0.33 | 3.61 ± 0.08 | 6.02 ± 0.03 | 6.47 ± 0.07 | 6.33 ± 0.47 | 6.36 ± 0.2 |
| 17 | G(anhM)4 + GM5 | 1.5 ± 0.08 | 1.57 ± 0.22 | 1.63 ± 0.13 | 2.3 ± 0.08 | 2.41 ± 0.06 | 2.15 ± 0.09 | 2.3 ± 0.1 |
| 18 | G(anhM)5 + GM4 | 1.94 ± 0.1 | 1.61 ± 0.08 | 1.49 ± 0.3 | 3.09 ± 0.04 | 3.07 ± 0.06 | 3.18 ± 0.08 | 3.12 ± 0.07 |
| 19 | G(anhM)3+ G(anhM)4 | 0.15 ± 0.02 | 0.15 ± 0.00 | 0.17 ± 0.03 | 0.46 ± 0.05 | 0.5 ± 0.05 | 0.41 ± 0.05 | 0.42 ± 0.06 |
| 20 | G(anhM)4+ G(anhM)4 | 0.41 ± 0.03 | 0.36 ± 0.04 | 0.47 ± 0.02 | 1.05 ± 0.07 | 1.22 ± 0.04 | 1.05 ± 0.08 | 1.15 ± 0.11 |
| 21 | G(anhM)5+ G(anhM)4 | 0.42 ± 0.04 | 0.45 ± 0.06 | 0.49 ± 0.06 | 0.93 ± 0.08 | 1.07 ± 0.02 | 0.84 ± 0.05 | 0.90 ± 0.05 |
| | *Additional muropeptides* | 0.42 ± 0.07 | 0.44 ± 0.03 | 0.44 ± 0.01 | 2.03 ± 0.14 | 3.11 ± 0.02 | 1.97 ± 0.23 | 1.71 ± 0.03 |
| 22 | GM4 + AmDap [c] | 0.15 ± 0.03 | 0.14 ± 0.01 | 0.14 ± 0.00 | 0.83 ± 0.07 | 1.29 ± 0.03 | 0.89 ± 0.10 | 0.73 ± 0.03 |
| 23 | GM4 + AmDap [d] | 0.06 ± 0.01 | 0.07 ± 0.00 | 0.07 ± 0.01 | 0.19 ± 0.01 | 0.28 ± 0.02 | 0.14 ± 0.05 | 0.16 ± 0.02 |
| 24 | GM4 + AAmDap [e] | 0.12 ± 0.02 | 0.12 ± 0.01 | 0.13 ± 0.01 | 0.64 ± 0.06 | 0.93 ± 0.01 | 0.64 ± 0.07 | 0.53 ± 0.02 |
| 25 | GM5 + AmDap [f] | 0.09 ± 0.02 | 0.12 ± 0.01 | 0.11 ± 0.01 | 0.37 ± 0.03 | 0.61 ± 0.02 | 0.30 ± 0.03 | 0.28 ± 0.01 |

GM2: GlcNAc-Mur*N*Ac-dipeptide; GM3: GlcNAc-Mur*N*Ac-tripeptide; GM4: GlcNAc-Mur*N*Ac-tetrapeptide; GM5: GlcNAc-Mur*N*Ac-pentapeptide; AnhM: *N*-acetyl-anhydromuramic acid.

[a] Percentages were calculated as in Glauner *et al.* (30).

[b] GM4+gly: GlcNAc-Mur*N*Ac-Tetrapeptide with an additional glycine moiety.

The structures of GM4 + AmDap [c], GM4 + AmDap [d], GM4 + AAmDap [e] and GM5 + AmDap [f] are represented below

(c)

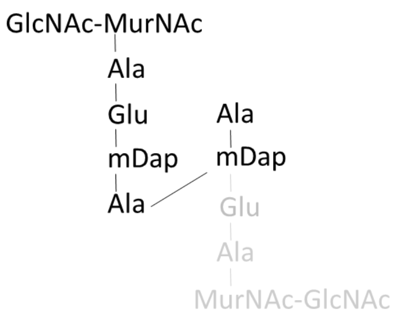
GlcNAc-MurNAc
Ala
Glu
mDap
Ala
Ala
mDap
Glu
Ala
MurNAc-GlcNAc

(d)

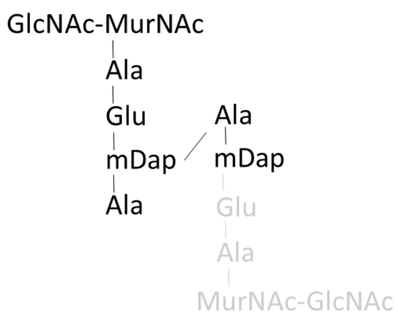
GlcNAc-MurNAc
Ala
Glu
mDap
Ala
Ala
mDap
Glu
Ala
MurNAc-GlcNAc

(e)

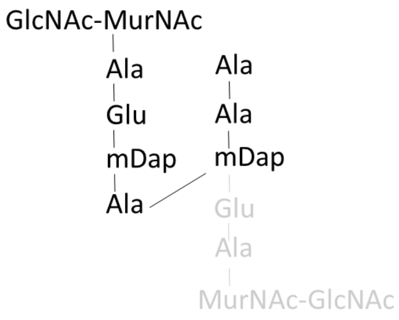
GlcNAc-MurNAc
Ala
Glu
mDap
Ala
Ala
Ala
mDap
Glu
Ala
MurNAc-GlcNAc

(f)

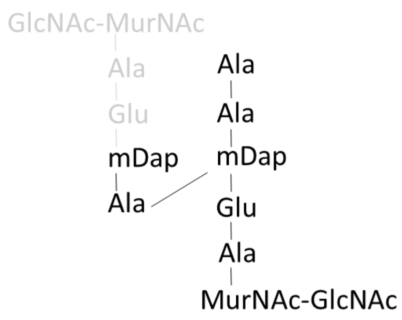
GlcNAc-MurNAc
Ala
Glu
mDap
Ala
Ala
Ala
mDap
Glu
Ala
MurNAc-GlcNAc

**Panel B:** Total ion current chromatograms of the HPLC/MS profiles of muropeptides of the seven samples indicated above (only one replicate is represented). Peak numbers correspond to those of Panel A and are identical in every graph.

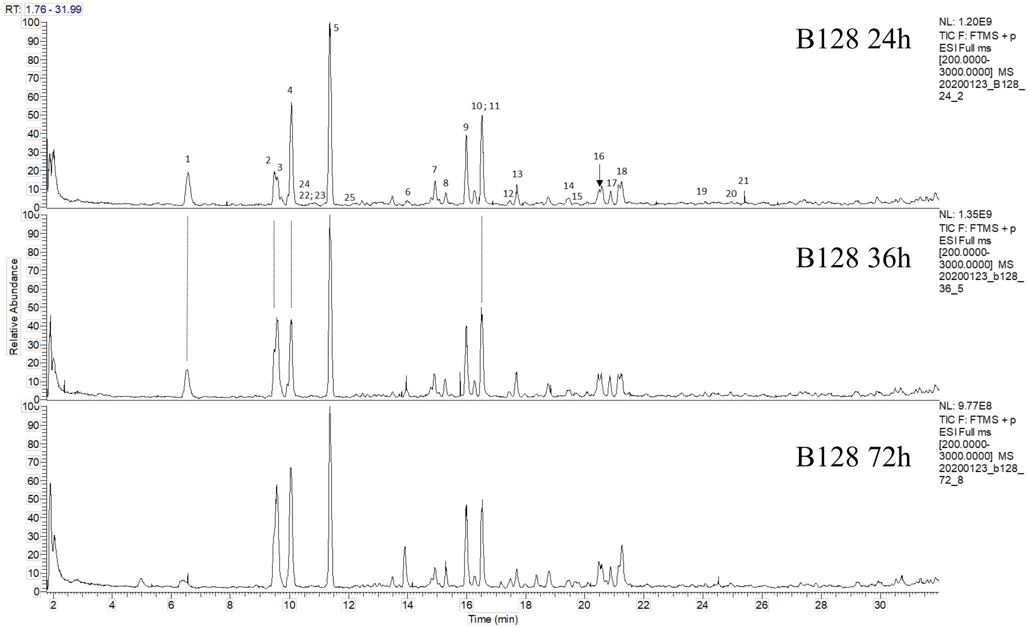


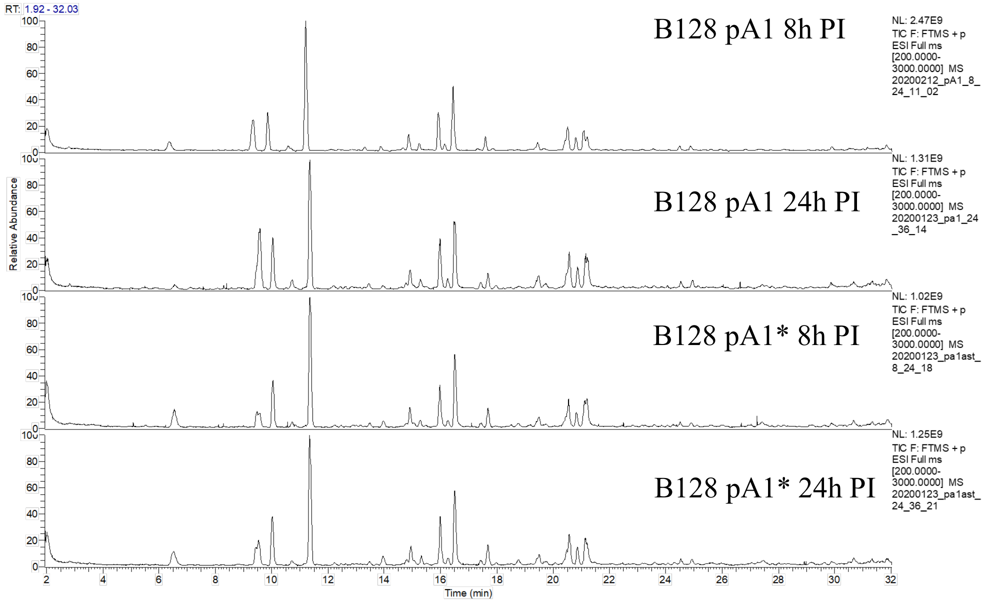

**Supplementary Table S2:** List of strains used in this study.

| | Strain designation | Genotype | Plasmid | Antibiotic resistance markers [a] | Reference or source |
|---|---|---|---|---|---|
| B128 | HPLEM001 | Parental wild type strain | - | - | (54, 55) |
| B128 Δ*rpsL::rpsL1* | HPLEM015 | Δ*rpsL::rpsL1* | - | Str | (31) |
| B128 Δ *TA A1::Kan + pA1-isoA1* | HPLEM011 | Δ*AapA1-isoA1::aphA-3* | pA1-isoA1 | Kan, Cm | (24) |
| B128 Δ *TA A1::Kan + pA1* | HPLEM012 | Δ*AapA1-isoA1::aphA-3* | pA1 | Kan, Cm | (24) |
| B128 Δ *TA A1::Kan + pA1** | HPLEM086 | Δ*AapA1-isoA1::aphA-3* | pA1* | Kan, Cm | (24) |
| B128 Δ*rpsL::rpsL1* Δ*A1 PureA-GFP* Δ*flaA::Apra + pA1* | HPLEM213 | Δ*rpsL::rpsL1* Δ*A1* Δ*flaA::Apra* Δ*ureA::GFP-mut2* | pA1 | Str, Kan, Apr, Cm | This study |
| B128 Δ*rpsL::rpsL1* Δ*TA A1 + pA1-GFP* | HPLEM155 | Δ*rpsL::rpsL1* Δ*A1* | pA1-GFP | Str, Cm | This study |
| B128 Δ*TA A1-isoA1:: AapA1-222nt-GFP-253nt* | HPLEM160 | Δ*A1-isoA1:: AapA1-222nt-gfp-mut2-253nt* | - | Kan | This study |
| B128 Δ *TA A1 AapA1-isoA1::PaapA1-lacZ-Kan* | HPLEM084 | Δ *AapA1-isoA1::PaapA1-lacZ-Kan* | - | Kan | This study |
| B128 Δ *TA A1 AapA1-isoA1::PisoA1-lacZ-Kan* | HPLEM142 | Δ *AapA1-isoA1::PisoA1-lacZ-Kan* | - | Kan | This study |
| B128 Δ*rpsL::rpsL1* Δ*TA* Δ*A5* Δ*A3* Δ*A1* Δ*A6* Δ*A4::Kan* | HPLEM159 | Δ*rpsL::rpsL1* Δ*TA* Δ*A5* Δ*A3* Δ*A1* Δ*A6* Δ*A4::Kan* | - | Str, Kan | This study |
| *B128* Δ*rpsL::L1* Δ*TA A1* | HPLEM214 | Δ*rpsL::L1* Δ*TA A1* | - | Str | This study |
| *B128* Δ*rpsL::L1* Δ*TA A3* | HPLEM080 | Δ*rpsL::L1* Δ*TA A3* | - | Str | This study |
| *B128* Δ*rpsL::L1* Δ*TA A4::Kan* | HPLEM157 | Δ*rps::L1* Δ*TA A4::Kan* | - | Str, Kan | This study |
| *B128* Δ*rpsL::L1* Δ*TA A5* | HPLEM067 | Δ*rpsL::L1* Δ*TA A5* | - | Str | This study |
| *B128* Δ*rps::L1* Δ*TA A6* | HPLEM216 | Δ*rps::L1* Δ*TA A6* | - | Str | This study |
| B128 Δ*rpsL::rpsL1* Δ*TA* Δ*A5* Δ*A3* Δ*A1* Δ*A6* Δ*A4::Kan* | HPLEM159 | Δ*rpsL::rpsL1* Δ*TA* Δ*A5* Δ*A3* Δ*A1* Δ*A6* Δ*A4::Kan* | - | Str, Kan | This study |

[a] Str: Streptomycin; Kan: Kanamycin; Apr: Apramycin; Cm: Chloramphenicol

Supplementary Table S3: List of plasmids used in this study.

| Name | Description | Resistance (*) | Reference or source |
|---|---|---|---|
| pILL2157 | Derivative of the pHeL-2 *E. coli-H. pylori* shuttle vector, carries *lacZ* under the control of p*ureI* with 2 LacI-binding sites | Cm | (27) |
| pILL2157bis | pILL2157 without *lacZ* (pILL2157bis) | Cm | This study |
| pDifWT-RC | *rpsL-cat* cassette flanked by *difH* | | (56) |
| pA1-isoA1 | *AapA1-isoA1* locus cloned into pILL2157bis | Cm | (24) |
| pA1 | *AapA1-isoA1* locus cloned into pILL2157bis with *isoA1* promoter inactivated | Cm | (24) |
| pA1* | *AapA1-isoA1* locus cloned into pILL2157bis with isoA1 promoter inactivated and AapA1 start codon mutated to ATT | Cm | (24) |
| pGEM ΔTA A1 | Suicide plasmid with *difH-rpsL-cat-difH* flanked by upstream and downstream region *A1* for markerless deletion of A1 locus. | Amp/Cm | This study |
| pGEM ΔTA A3 | Suicide plasmid with *difH-rpsL-cat-difH* flanked by upstream and downstream region *A3* for markerless deletion of A3 locus. | Amp/Cm | This study |
| pGEM ΔTA A5 | Suicide plasmid with *difH-rpsL-cat-difH* flanked by upstream and downstream region *A5* for markerless deletion of A5 locus. | Amp/Cm | This study |
| pGEM ΔTA A6 | Suicide plasmid with *difH-rpsL-cat-difH* flanked by upstream and downstream region *A6* for markerless deletion of A6 locus. | Amp/Cm | This study |
| pJET-pureA-GFP-mut2-Kan-ureA | Suicide plasmid with *gfp-mut2* under the control of *ureA* promoter for chromosomal integration at the *ureA* locus | Amp/Kan | (61) |
| pA1-GFP | *gfp-mut2* cloned in translational fusion between AapA1 sequence 222 nt (from 0 to the 222 nt aapA1) and 253 nt (from 223 to the 253 nt) into vector pILL2157bis. AapA1-222nt-GFP-253nt is under the control of the *ureI* promoter. | Cm | This study |

(*) Cm: Chloramphenicol; Amp: Ampicillin; Kan: Kanamycin

Supplementary Table S4: list of primers used in this study

| Name | Sequence 5'→3' | Description |
|---|---|---|
| **oLEM001** | GTAAGCATTGCCGACAAACAC | ***rpsl::rpsl1*** Forward primer to amplify upstream region of *rpsL.* |
| **oLEM002** | CCAATTGATTTATGGTAGGCACTATTTTTCCTTATTC | ***rpsl::rpsl1*** Reverse primer to amplify upstream region of *rpsL.* Primer contains a homolougous region to *rpsL1.* |
| **oLEM003** | GTGCCTACCATAAATCAATTGG | ***rpsl::rpsl1*** Forward primer to amplify *rpsL1* from pDifWT-RC. |
| **oLEM004** | CTAACGGATTTGTCTGTATG | ***rpsl::rpsl1*** Reverse primer to amplify *rpsL1* from pDifWT-RC. |
| **oLEM005** | CATACAGACAAATCCGTTAGAGGAAAACAAAAACATGAGAAG | ***rpsl::rpsl1*** Forward primer to amplify downstream region of rpsL. Primer contains a homologous region to *rpsL1.* |
| **oLEM006** | CCATTCTAACTCCAATTACCAG | ***rpsl::rpsl1*** Reverse primer to amplify downstream region of *rpsL.* |
| **oLEM192** | GGATGTATAGACCGTTATGG | ***flaA::apr*** Forward primer to amplify upstream region of *flaA.* |
| **oLEM193** | cactccCTAgTTAgTCACcatTGTTGTAACTCCTTG | ***flaA::apr*** Reverse primer to amplify upstream region of *flaA*. Primer contains a homolougous region to *apr* resistance cassette with a stop codon and an RBS. |
| **oLEM120** | TGAcTAAcTAGggagtgcaATGtcgtgcaa | ***flaA::apr*** Forward primer to amplify *apr* resistance cassette with stop codon upstream of the RBS. |
| **oLEM066** | cgatccgctccacgtgttgcc | ***flaA::apr*** Reverse primer to amplify *apr* resistance cassette. |
| **oLEM194** | ggcaacacgtggagcggatcgCAAGCCAATACCGTTCAAC | ***flaA::apr*** Forward primer to amplify downstream region of *flaA*. Primer contains a homologous region to *apr* resistance cassette. |
| **oLEM195** | CATAGCATAAAATCGCATCC | ***flaA::apr*** Reverse primer to amplify downstream region of *flaA*. |
| **oLEM015** | CTCCCACCGCAATTGATTG | **Δ*A1* with marker less system** Forward primer to amplify upstream region of *A1* Toxin antitoxin locus. |
| **oLEM035** | CATACTCGAGGCTTGATTGAGTGCATCAAAAC | **Δ*A1* with marker less system** Reverse primer to amplify upstream region of *A1* Toxin antitoxin locus. Primer contains a *Xho*I restriction site. |
| **oLEM036** | CATAGGATCCCGAAGTTTCTGTAAAACGATAG | **Δ*A1* with marker less system** Forward primer to amplify downstream region of A1 Toxin antitoxin locus. Primer contains a *Bam*HI restriction site. |
| **oLEM018** | CTCAATGCGTTTAGGATTAATC | **Δ*A1* with marker less system** Reverse primer to amplify downstream region of *A1* Toxin antitoxin locus. |
| **oLEM019** | CATTCAAAGATGTTGGTAG | **Δ*A3* with marker less system** Forward primer to amplify upstream region of *A3* Toxin antitoxin locus. |
| **oLEM037** | CATACTCGAGCTAGATCGCATCCAATACG | **Δ*A3* with marker less system** Reverse primer to amplify upstream region of *A3* Toxin antitoxin locus. Primer contains a *Xho*I restriction site. |
| **oLEM038** | CATAGGATCCCAAGAGCGTTCCCTTAAGC | **Δ*A3* with marker less system** Forward primer to amplify downstream region of *A3* Toxin antitoxin locus. Primer contains a *Bam*HI restriction site. |

| | | |
|---|---|---|
| **oLEM022** | CTTGAAAGGCTTCAATCAAG | **Δ*A3* with marker less system**<br>Reverse primer to amplify downstream region of *A3* Toxin antitoxin locus. |
| **oLEM027** | CATGCTTGTCAAACCACAG | **Δ*A5* with marker less system**<br>Forward primer to amplify upstream region of *A5* Toxin antitoxin locus. |
| **oLEM041** | CATACTCGAGGCTCTTAAATGCAACCAC | **Δ*A5* with marker less system**<br>Reverse primer to amplify upstream region of *A5* Toxin antitoxin locus. Primer contains a *Xho*I restriction site. |
| **oLEM042** | CATAGGATCCCTAAGAGCGTTCCCCTAAG | **Δ*A5* with marker less system**<br>Forward primer to amplify downstream region of *A5* Toxin antitoxin locus. Primer contains a *Bam*HI restriction site. |
| **oLEM030** | CTCAGTATGTGAATTTAGCG | **Δ*A5* with marker less system**<br>Reverse primer to amplify downstream region of *A5* Toxin antitoxin locus. |
| **oLEM031** | GCCAAGCACCATCTTCTTTATG | **Δ*A6* with marker less system**<br>Forward primer to amplify upstream region of *A6* Toxin antitoxin locus |
| **oLEM043** | CATACTCGAGGCTGCAAACCACTCATTTAAAG | **Δ*A6* with marker less system**<br>Reverse primer to amplify upstream region of *A6* Toxin antitoxin locus. Primer contains a *Xho*I restriction site. |
| **oLEM044** | CATAGGATCCGGGTTATCCTTAAGTGGA | **Δ*A6* with marker less system**<br>Forward primer to amplify downstream region of *A6* Toxin antitoxin locus. Primer contains a *Bam*HI restriction site |
| **oLEM034** | CTCATTACGACACTATTGC | **Δ*A6* with marker less system**<br>Reverse primer to amplify downstream region of *A6* Toxin antitoxin locus. |
| **oLEM045** | CATACTCGAGatttaaaagtttgaaaagtgcag | **marker less system**<br>Forward primer to amplify *rpsl-cat* cassette from pDifWT-RC. Primer contains a *Xho*I restriction site. |
| **oLEM046** | CATAGGATCCtcatttagttatgaaaactgcac | **marker less system**<br>Reverse primer to amplify *rpsl-cat* cassette from pDifWT-RC. Primer contains a *Bam*HI restriction site. |
| **oLEM023** | GAGGCTGTAAGGATAAGG | **Δ*A4::Kan***<br>Forward primer to amplify upstream region of *A4* Toxin antitoxin locus. |
| **oLEM107** | gTTAgTCAcccgggtaccCAAACGCTAAAACGAGGCAC | **Δ*A4::Kan***<br>Reverse primer to amplify upstream region of *A4* Toxin antitoxin locus. Primer contains a homologous region to the Kanamycin resistance cassette. |
| **oLEM009** | GGTACCCGGGTGACTAAC | **Δ*A4::Kan***<br>Forward primer to amplify the Kanamycin resistance cassette. |
| **oLEM010** | CATTATTCCCTCCAGGTAC | **Δ*A4::Kan***<br>Reverse primer to amplify the Kanamycin resistance cassette |
| **oLEM108** | gtacctggagggaataATGGTTGGTCATTTTGGTATAAAAC | **Δ*A4::Kan***<br>Forward primer to amplify downstream region of *A4* Toxin antitoxin locus. Primer contains a homologous region to the Kanamycin resistance cassette. |
| **oLEM026** | CCCTAATAGTAGAAAATGGAG | **Δ*A4::Kan***<br>Reverse primer to amplify downstream region of *A4* Toxin antitoxin locus. |
| **oLEM073** | GATCATTAAAGGCTCCTTTTG | ***pA1-GFP***<br>Forward primer to amplify upstream region of purel in pILL2157bis. |
| **oLEM080** | CAAAATGCCCGCTTCAATAAAAC | ***pA1-GFP***<br>Reverse primer to amplify *aapA1* where stop codon of the A1 toxin has been replaced by Ala codon . |

| Name | Sequence 5'→3' | Description |
|---|---|---|
| **oLEM081** | GAAGCGGGCATTTTGTAAAACGAAGTTTCTGGCAAACGATAG | ***pA1-GFP*** Forward primer to amplify 3'-UTR of aapA1 (from 142 to 222 nt). Contains a homologous region to oLEM080 and 2 stop codons each mutated in Ala codon to enable the GFP translational fusion. |
| **oLEM083** | GTTCTTCTCCTTTACTCATGAAAACCCTTAAAGCTAAAAG | ***pA1-GFP*** Reverse primer to amplify 3'-UTR region of *aapA1*. Primer also contains a homologous region to *gfp-mut2.* |
| **oLEM082** | ATGAGTAAAGGAGAAGAAC | ***pA1-GFP*** Forward primer to amplify *gfp-mut2.* |
| **oLEM084** | GTCATTTGTATAGTTCATCC | ***pA1-GFP*** Reverse primer to amplify *gfp-mut2.* |
| **oLEM085** | GGATGAACTATACAAATGACTTTAAAGCTTATCCTTAAC | ***pA1-GFP*** Forward primer to amplify *aapA1* 3'-UTR (from 222 nt to 253 nt). Primer contains a homologous region to *gfp-mut2.* |
| **oLEM099** | CGGTACCCTAGTATTCCAAGCAAAGAATG | ***pA1-GFP*** Reverse primer to amplify *aapA1* 3'-UTR from pA1. Primer contains a KpnI restriction site for cloning in pILL2157bis. |
| **oLEM047** | atgACTAGTACCATGATTACG | ***lacZ transcriptional fusion*** Forward primer to amplify *lacZ* from pILL2157bis. |
| **oLEM069** | GAAGGAAAAGcatatgACTAG | ***lacZ transcriptional fusion*** Forward primer to amplify *lacZ* with RBS from pILL2157bis |
| **oLEM048** | GGGATCCTTATTTTTGACAC | ***lacZ transcriptional fusion*** Reverse primer to amplify *lacZ* from pILL2157bis. |
| **oLEM059** | GTGTCAAAAATAAGGATCCCGGTACCCGGGTGACTAAC | ***lacZ-Kan*** Forward primer to amplify Kan cassette with oLEM010. Primer contains a homologous region to *lacZ*. |
| **oLEM049** | CGTAATCATGGTACTAGTcatGACAAACTCCTTTTATGG | ***PaapA1-lacZ*** Reverse primer to amplify upstream region of *aapA1* with oLEM015. Primer contains a homologous region to *lacZ*. |
| **oLEM060** | gtacctggagggaataATGCGAAGTTTCTGTAAAACGATAG | ***PaapA1-lacZ*** Forward primer to amplify downstream region of *aapA1* with oLEM018. Primer contains a homologous region to *Kan*. |
| **oLEM070** | CTAGTcatatgCTTTTCCTTCAACATGGCAAAAACTCTTGG | ***PisoA1-lacZ*** Reverse primer to amplify the upstream region of isoA1 promoter with oLEM018. Primer contains a homologous region to RBS-*lacZ*. |
| **oLEM071** | gtacctggagggaataATGCAACAATCTTTCTAAAACC | ***PisoA1-lacZ*** Forward primer to amplify downstream region of isoA1with oLEM015. Primer contains a homologous region to Kanamycin resistance cassette. |
| **oLEM213** | AGTTTTTGCCATGTTTGGTA | ***Northern Blot*** *AapA1 probe* |
| **oLEM214** | GAGTTTGTCATGGCTACCAA | ***Northern Blot*** *isoA1 probe* |
| **oLEM215** | TCGGAATGGTTAACTGGGTAGTTCCT | ***Northern Blot*** *5S probe* |
