## Supplementary Figures S1-S2-S3-S4-S6-S7-S8-S9 for "A peptide of a type I toxin-antitoxin system induces *Helicobacter pylori* morphological transformation from spiral-shape to coccoids"

### Legends of supplementary figures

**Fig. S1: Schematic representation of the native A1 locus, of the AapA1-GFP fusions expressed from plasmid pILL2157 or from the native locus and of the P*aapA1-lacZ* and P*IsoA1-lacZ* fusions expressed at their native loci from their respective native promoters**

SD indicates the Shine-Dalgarno sequence; -10, the position of the -10 box of the promoter; PureI, the promoter of the *ureI* gene from plasmid pILL2157; arrows, the direction of transcription; a red cross on an arrow, an inactivated promoter; GFP, green fluorescent protein; kan, the gene conferring kanamycin resistance; a star (*) a STOP codon that has been replaced by a codon coding for Ala. A hatched *IsoA1* arrow indicates that the corresponding RNA is not expressed. The corresponding strains are listed in Table S2, the plasmids in Table S3 and the primers used for the construction in Table S4.

**Fig. S2: Controls of the *H. pylori* fractionation procedure**

**A.** Western blot analysis of total extract (T), soluble extract (SE), inner membrane (IM) and outer membrane (OM) fractions prepared from a control *H. pylori* B128 strain and revealed with the following control antibodies anti-Pbp2 for IM, anti-AlpA for the OM and anti-AmiE for the cytoplasmic fraction.

**B**. Fluorescence of a strain expressing GFP from plasmid pILL2157 is presented as a control for the data of **Fig. 2B**.

**Fig. S3: Representative flow cytometry histograms of different *H. pylori* cells stained with a PMF-sensitive dye**

Histograms of fluorescence intensity of cells from strains pA1-isoA1, pA1 or pA1* stained with the pmf-sensitive MitoTracker Red CMXROS dye at 0h (**A**), 8h (**B**) or 24h (**C**) after 1 mM IPTG addition or not. pA1* cells treated with TCS served as a control for PMF dissipation conditions.

**Fig S4: Measurement of *H. pylori* cell length during growth over time**

Mean curve obtained from the analysis of the growth of 61 individual *H. pylori* bacteria (strain HPLEM213 without IPTG, **Table S2**). Bacteria were analyzed by live microscopy, and their size and time of division were measured from their separation following a division to the next division. The curves were analyzed and normalized. Mean division time is 165 min, initial length mean is 1.9 µm, mean length when division occurs is 3.2 µm.

**Fig. S5: Movies of *H. pylori* morphological transformation**

Representative movies of the transformation of *H. pylori* upon expression of AapA1 toxin. Snapshots of the cells were taken at intervals of 10min during 10h.

**Fig. S6: Response of the *aapA1* and *IsoA1* promoters to different stresses**

β-galactosidase activities expressed by strains expressing the P*aapA1-lacZ* and P*IsoA1-lacZ* fusions from the native locus were measured after 6h treatment with different stresses, $NiCl_2$ (20 and 200 mM), pH 4, Tetracycline (0.1 and 1 mg/ml) or Rifampicin (0.05 and 0.5 mg/ml). β-galactosidase activities are presented as ratio (expressed in %) of activities measured with stress versus activities of untreated samples. Results from 3 independent experiments performed in duplicates are shown. Error bars represent the standard deviation, NS corresponds to non-significant, (*P* >0.05).

**Fig. S7: Half-life of the *aapA1* and *IsoA1* RNAs**

RNA decay was determined by plotting normalized intensities (RNA signal relative to time 0) of bands corresponding to full length *aapA1* and *IsoA1* transcripts as a function of time after rifampicin addition. Approximate half-lives (min) measurements from three independent experiments are indicated for each transcript.

**Fig. S8: Growth, viability and morphological transformation of B128 WT strain and multiple isogenic class A TA systems deletion mutants under normal conditions or upon exposure to hydrogen peroxide**

**A)** Quantification of the proportion of coccoids and spiral bacteria in strain B128 and the Δ5 mutant at 6 and 24h without treatment or post-treatment with 2 or 5% $H_2O_2$. The images were analyzed with the MicrobeJ plugin from ImageJ.

**B)** Growth of the B128 WT strain, of six isogenic mutants carrying *AapA1-IsoA1* deletions (*ΔA1, ΔA2, ΔA3, ΔA4-2+A4-2, ΔA5 or ΔA6*) and of a multiple mutant strain carrying deletions of every functional class A TA system (Δ*5*) was followed under normal conditions during 44h. The growth curve of the mutants was similar to that of the parental WT strain.

**C)** Viability of the B128 WT strain and of the Δ5 multiple TA mutant was measured during 72h by determining colony forming units (CFU) by plating on blood agar medium. No significant difference was observed in the kinetics of loss of viability between these strains.

**D)** Exponentially growing B128 WT strain, Δ*A1* and Δ*5* isogenic mutants were exposed to 1% hydrogen peroxide. Their viability was measured during 12h by counting the colony forming units (CFU/mL) on blood agar plates.

**E)** Exponentially growing B128 WT strain, Δ*A1* and Δ*5* isogenic mutants were exposed to 1% hydrogen peroxide during 8 h. The percentage of survival was calculated by dividing the number of CFU/mL in the culture after 8h with hydrogen peroxide by the number of CFU/mL after 8 h of incubation without stress.

**Fig. S9: Sequence alignment of the six functional class A TA modules present on the *H. pylori* B128 genome**

Sequences alignment of the six functional class A TA module colored according to the percentage identity with their consensus sequence. The -10 box, Shine-Dalgarno (SD) and AapA toxin coding sequence are framed in red and the start and stop codons are indicated above the sequences. *IsoA* antisense RNA sequence is represented by a green bar under the sequence alignment. Note that in B128, two consecutive TA modules are found at the locus A4, here referred as locus A4-1 and locus A4-2. A locus corresponding to the position of the A2 locus in other *H. pylori* strains was identified but its corresponding A2 ORF was inactivated in B128 strain.

**Figure S1**

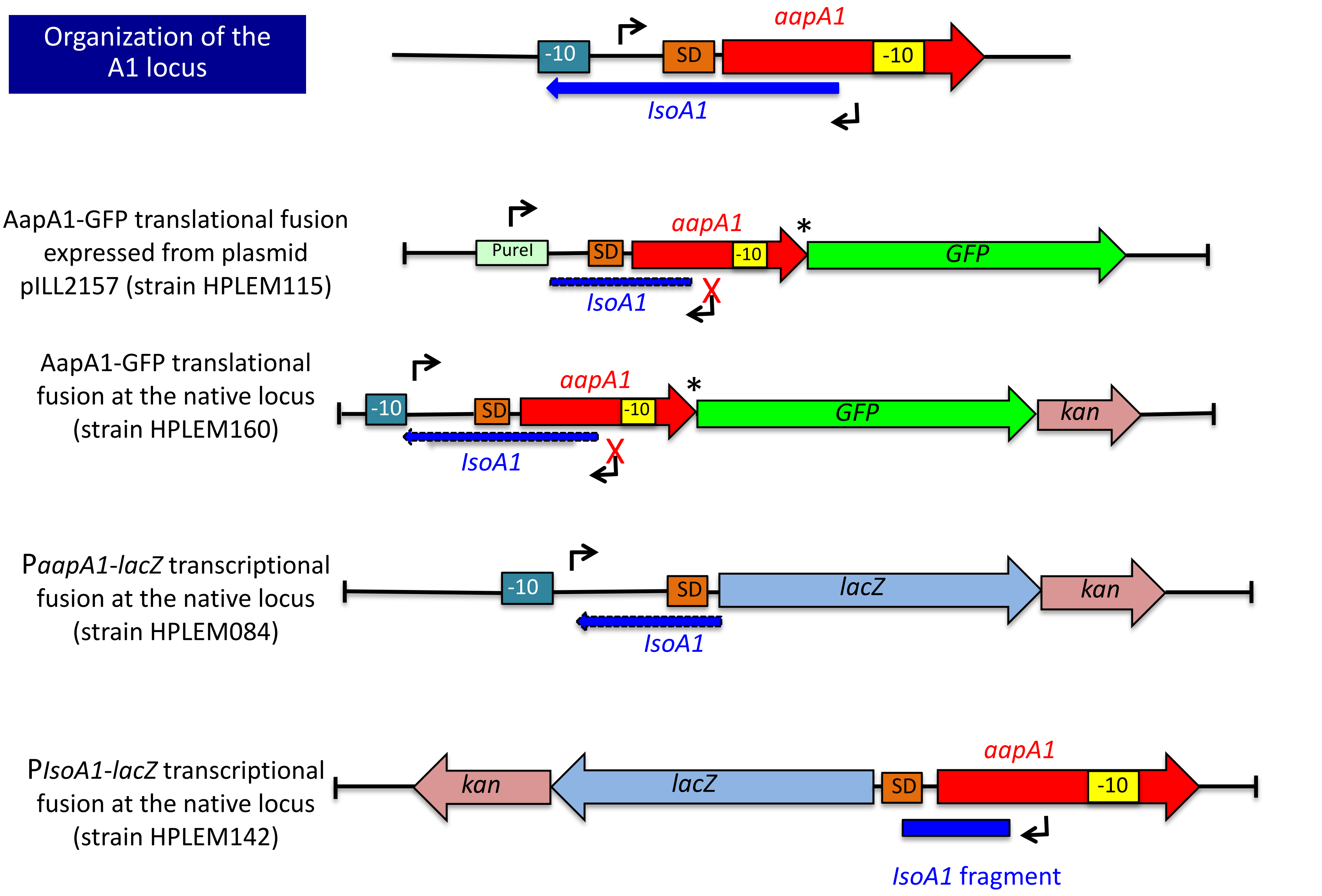

Organization of the A1 locus
aapA1
-10
SD
-10
IsoA1
AapA1-GFP translational fusion expressed from plasmid pILL2157 (strain HPLEM115)
PureI
SD
aapA1
-10
*
GFP
IsoA1
X
AapA1-GFP translational fusion at the native locus (strain HPLEM160)
-10
SD
aapA1
-10
*
GFP
kan
IsoA1
X
PaapA1-lacZ transcriptional fusion at the native locus (strain HPLEM084)
-10
SD
lacZ
kan
IsoA1
PIsoA1-lacZ transcriptional fusion at the native locus (strain HPLEM142)
kan
lacZ
SD
aapA1
-10
IsoA1 fragment

Figure S2

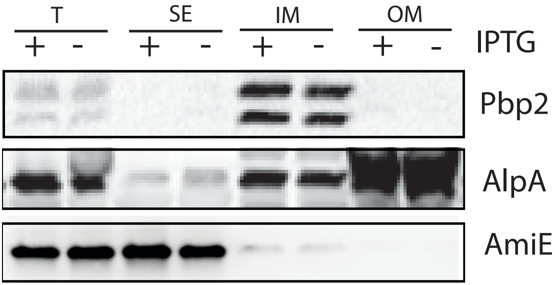


B

B128 GFP

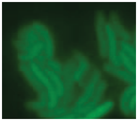

### Figure S3

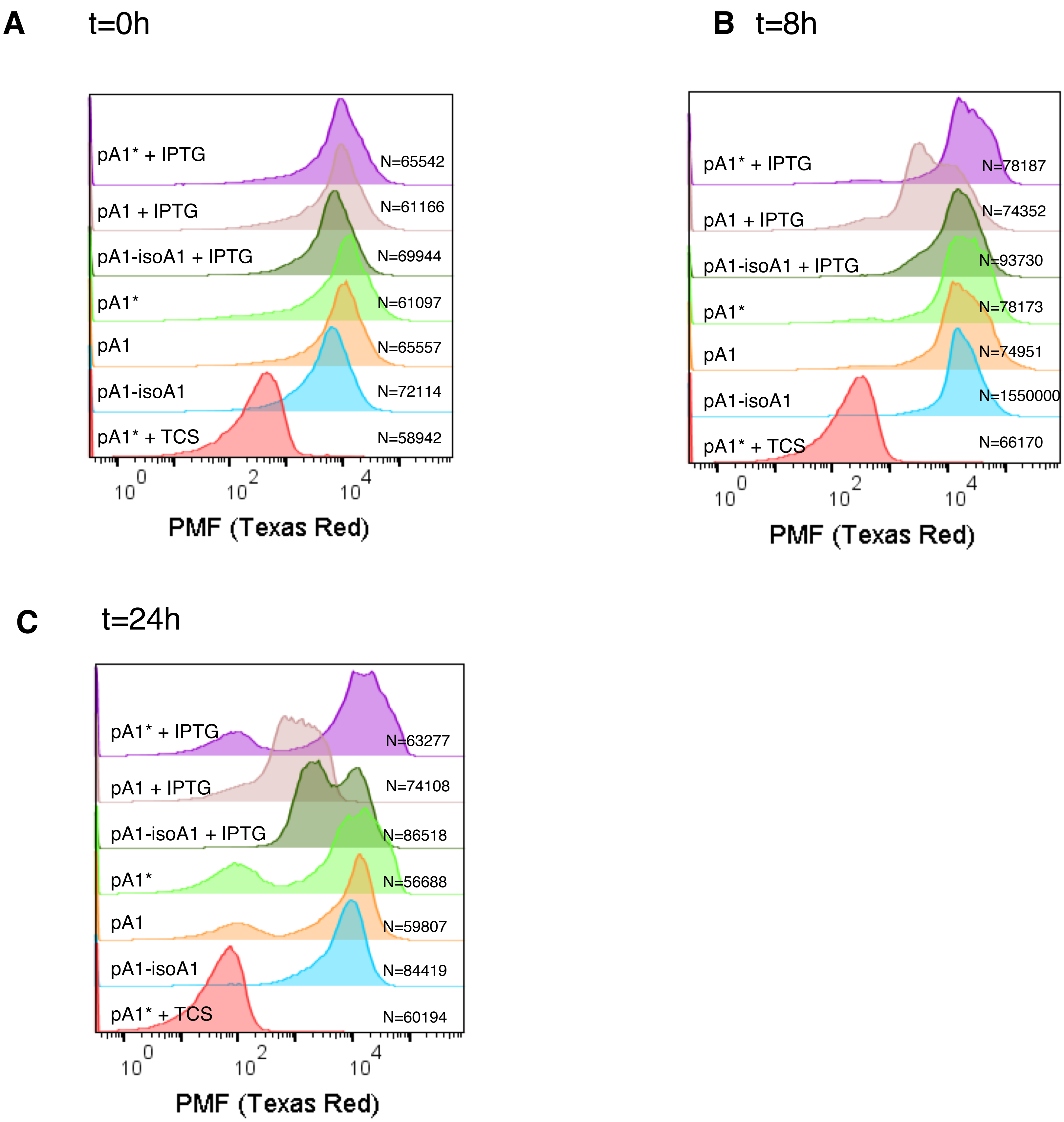
A
t=0h
pA1* + IPTG
N=65542
pA1 + IPTG
N=61166
pA1-isoA1 + IPTG
N=69944
pA1*
N=61097
pA1
N=65557
pA1-isoA1
N=72114
pA1* + TCS
N=58942
10^0
10^2
10^4
PMF (Texas Red)
B
t=8h
pA1* + IPTG
N=78187
pA1 + IPTG
N=74352
pA1-isoA1 + IPTG
N=93730
pA1*
N=78173
pA1
N=74951
pA1-isoA1
N=1550000
pA1* + TCS
N=66170
10^0
10^2
10^4
PMF (Texas Red)
C
t=24h
pA1* + IPTG
N=63277
pA1 + IPTG
N=74108
pA1-isoA1 + IPTG
N=86518
pA1*
N=56688
pA1
N=59807
pA1-isoA1
N=84419
pA1* + TCS
N=60194
10^0
10^2
10^4
PMF (Texas Red)

**Figure S4**

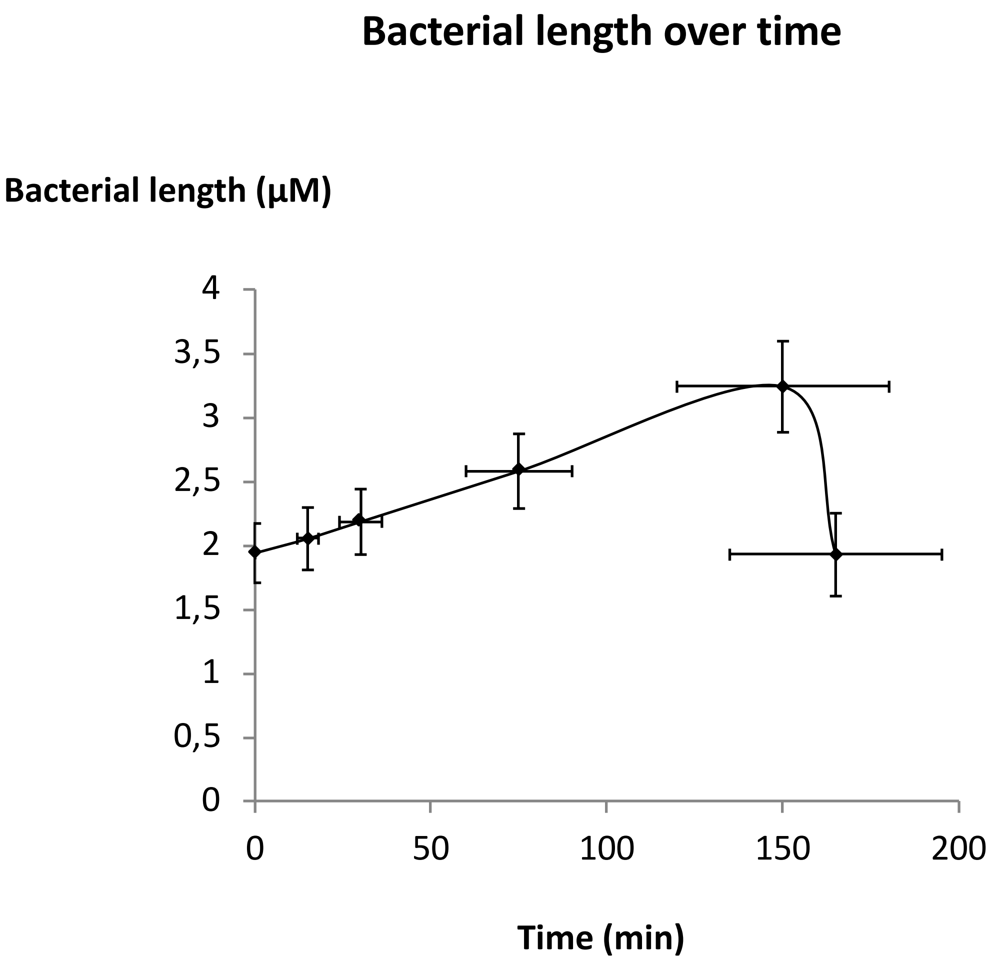
Bacterial length over time
Bacterial length (µM)
4
3,5
3
2,5
2
1,5
1
0,5
0
0
50
100
150
200
Time (min)

Figure S6

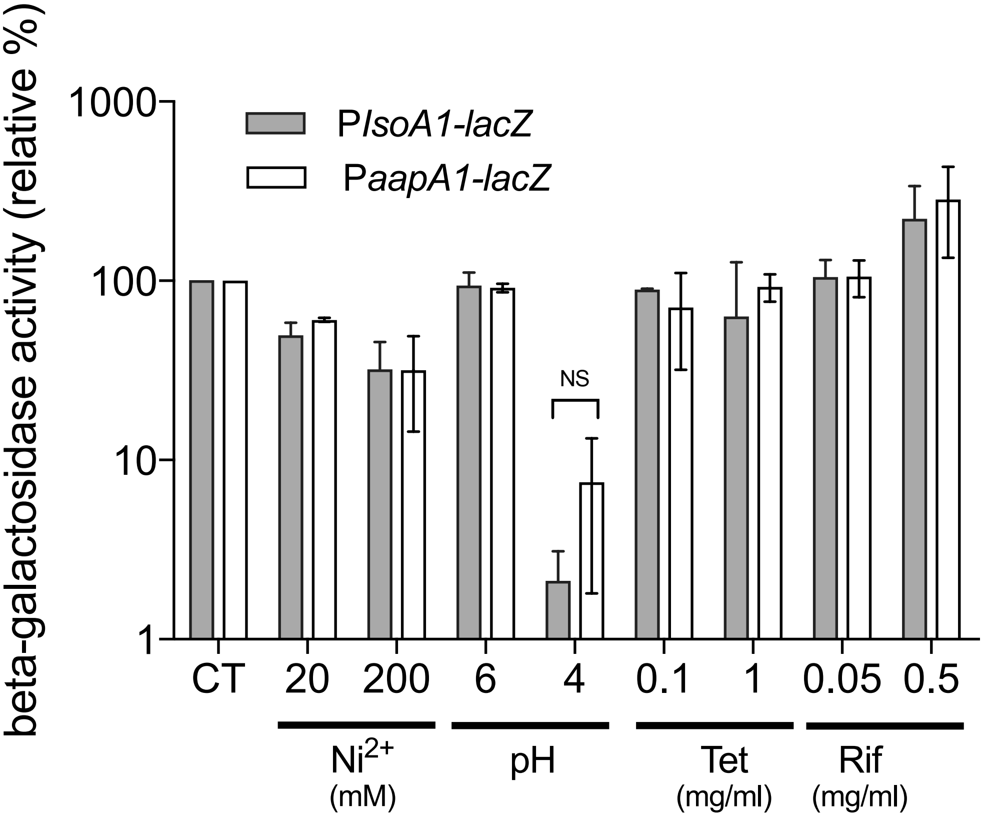
beta-galactosidase activity (relative %)
1000
100
10
1
PlsoA1-lacZ
PaapA1-lacZ
NS
CT
20
200
6
4
0.1
1
0.05
0.5
Ni2+
(mM)
pH
Tet
(mg/ml)
Rif
(mg/ml)

**Figure S7**

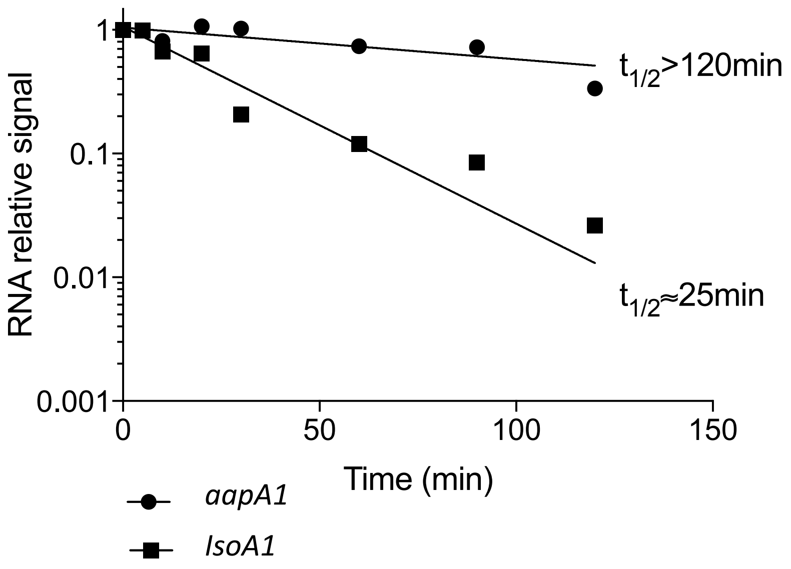
RNA relative signal
1
0.1
0.01
0.001
0
50
100
150
Time (min)
t1/2>120min
t1/2≈25min
aapA1
IsoA1

Figure S8

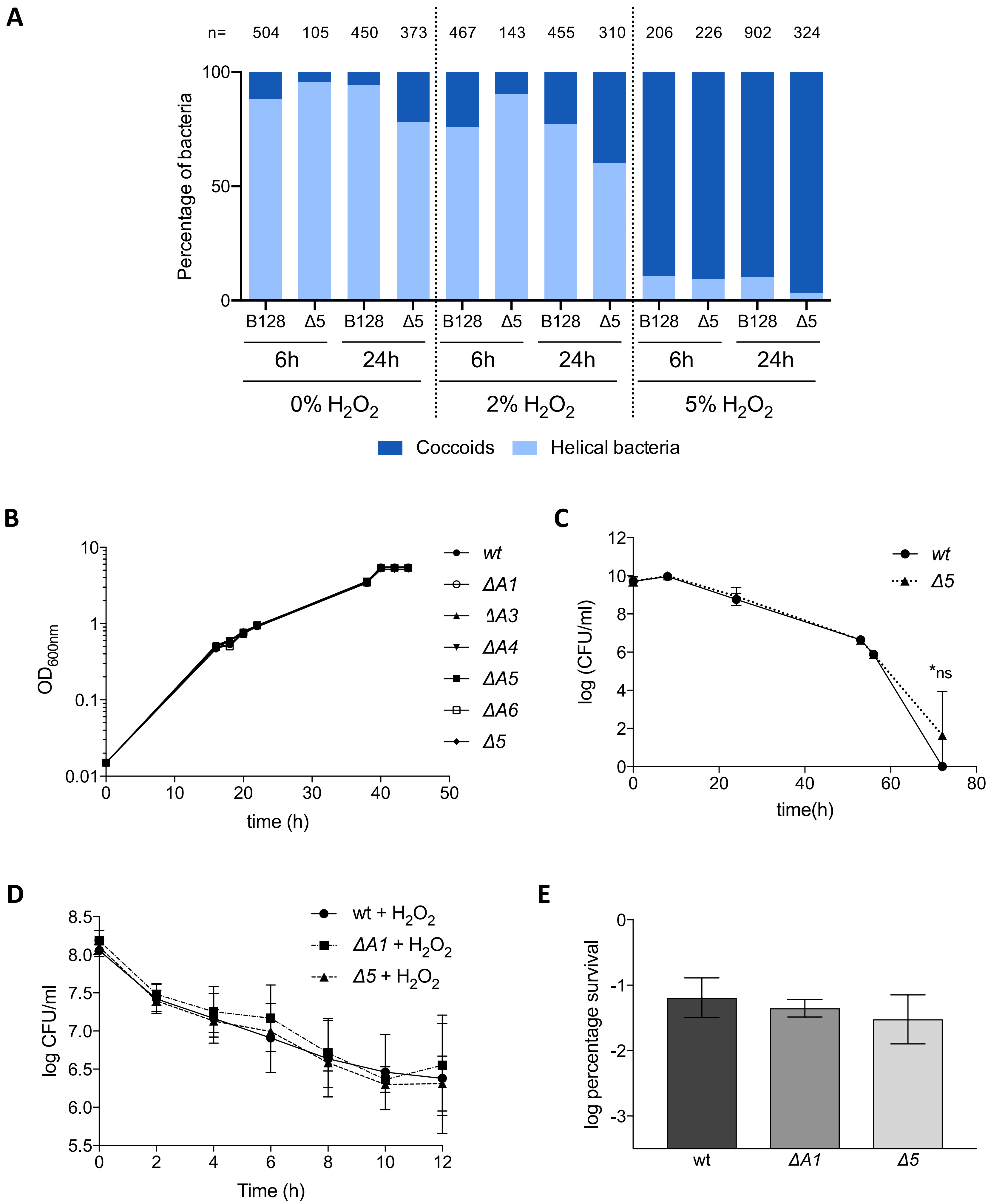
A
n= 504 105 450 373 467 143 455 310 206 226 902 324
Percentage of bacteria
100
50
0
B128 Δ5 B128 Δ5 B128 Δ5 B128 Δ5 B128 Δ5 B128 Δ5
6h 24h 6h 24h 6h 24h
0% H2O2
2% H2O2
5% H2O2
Coccoids
Helical bacteria
B
OD600nm
10
1
0.1
0.01
0 10 20 30 40 50
time (h)
wt
ΔA1
ΔA3
ΔA4
ΔA5
ΔA6
Δ5
C
log (CFU/ml)
12 10 8 6 4 2 0
0 20 40 60 80
time(h)
wt
Δ5
*ns
D
log CFU/ml
8.5 8.0 7.5 7.0 6.5 6.0 5.5
0 2 4 6 8 10 12
Time (h)
wt + H2O2
ΔA1 + H2O2
Δ5 + H2O2
E
log percentage survival
0 -1 -2 -3
wt
ΔA1
Δ5

Figure S9

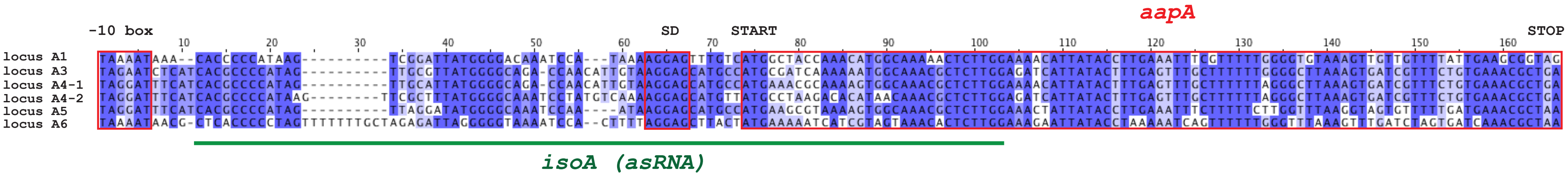
aapA
-10 box
SD
START
STOP
10 20 30 40 50 60 70 80 90 100 110 120 130 140 150 160
locus A1 TAAAATAAA--CACCCCCATAAG----------TCGGATTATGGGGACAAATCCA---TAAAAGGAGTTTGTCATGGCTACCAAACATGGCAAAAACTCTTGGAAAACATTATACCTTGAAATTTCGTTTTTGGGGTGTAAAGTTGTTGTTTTATTGAAGCGGTAG
locus A3 TAGAATCTCATCACGCCCCATAG----------TTGCGTTATGGGGCAGA-CCAACATTGTAAGGAGCATGCCATGCGATCAAAAAATGGCAAACGCTCTTGGAGATCATTATACTTTGAGTTTGCTTTTTTGGGGCTTAAAGTGATCGTTTCTGTGAAACGCTGA
locus A4-1 TAGGATTTCATCACGCCCCATAG----------TTGCATTATGGGGCAGA-CCAACATTGTAAGGAGCATGCCATGAAACGCAAAAGTGGCAAACGCTCTTGGAAAACATTATACTTTGAGTTTGCTTTTTTAGGGCTTAAAGTGATCGTTTCTGTGAAACGCTGA
locus A4-2 TAGGATTTCATCACGCCCCATAAG---------TTCGCTTTATGGGGCAAATCCTATGTCAAAAGGAGCATGTTATGCCTAAGACACATAACAAACGCTCTTGGAGATCATTATACTTTGAGTTTGCTTTTTTAGGGCTTAAAGTGATCGTTTCTGTGAAACGCTGA
locus A5 TAGGATTTCATCACGCCCCATAG----------TTAGGATATGGGGCAAATCCAA----ATAAGGAGCATGCCATGAAGCGTAAAAGTGGCAAACGCTCTTGGAAACTATTATACCTTGAAATTTCTTTTTTCTTGGTTTAAGGTAGTGTTTTTGATGAAACGCTAA
locus A6 TAAAATAACG-CTCACCCCCTAGTTTTTTTGCTAGAGATTAGGGGGTAAAATCCA--CTTTTAGGAGCTTACTATGAAAAATCATCGTAGTAAACACTCTTGGAAAGAATTATACCTAAAAATCAGTTTTTTGGGTTTAAAGTTTGATCTAGTGATCAAACGCTAA
isoA (asRNA)
