## Supplementary figures and images for "A peptide of a type I toxin-antitoxin system induces *Helicobacter pylori* morphological transformation from spiral-shape to coccoids"

### Fig. S5.pptx

#### Slide 1
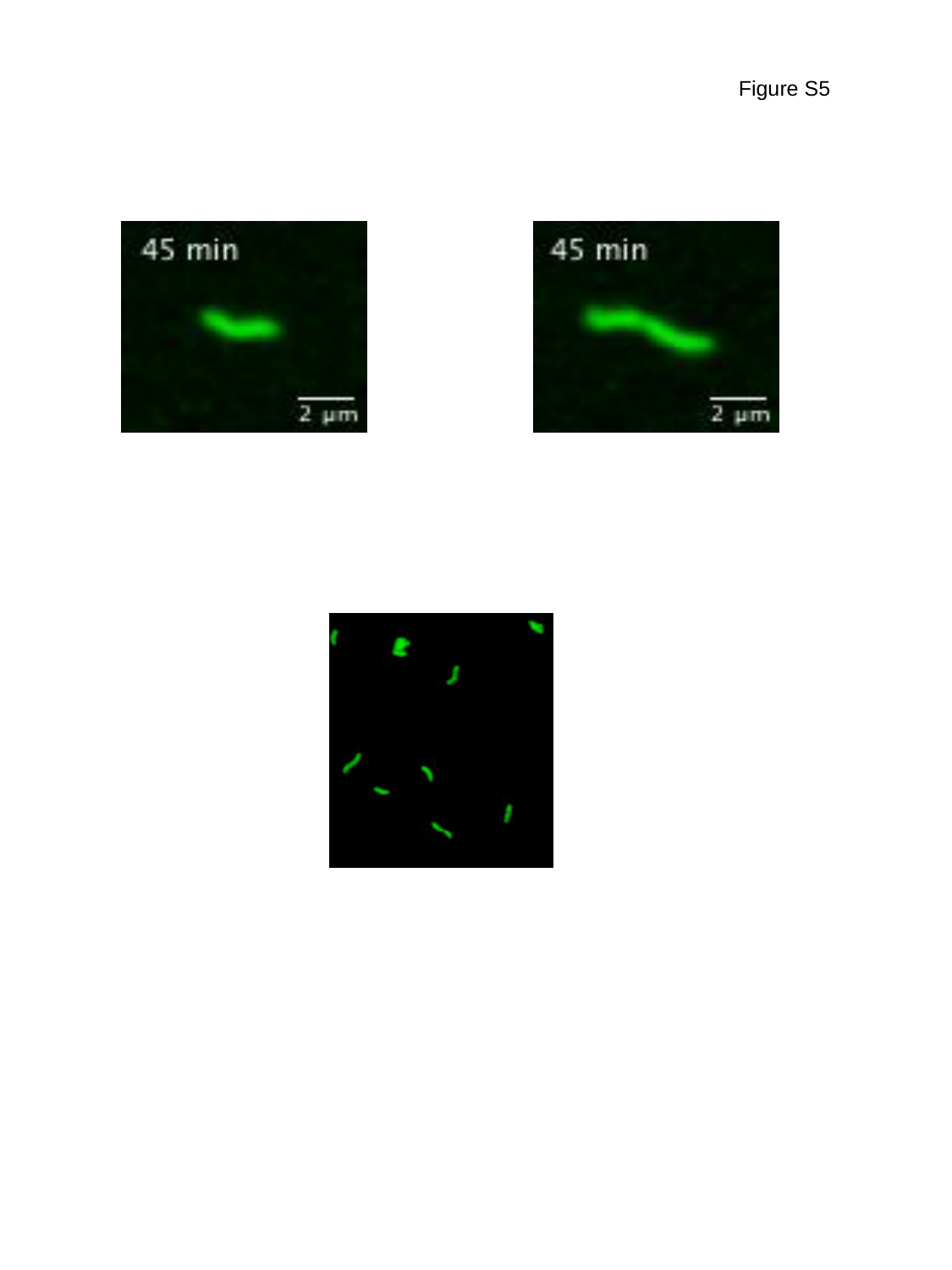

Figure S5
